# Three-dimensional topological defects and quasi-long-range order in biological liquid crystals

**DOI:** 10.1101/2025.04.14.648711

**Authors:** Anna E. Argento, Maria L. Varela, Gurveer Singh, Daiana P. Visnuk, Binyamin Jacobovitz, Mary E. Rutherford, Marta B. Edwards, Quentin Chaboche, Daniel A. Orringer, Jason A. Heth, Maria G. Castro, Daniel A. Beller, Carles Blanch-Mercader, Pedro R. Lowenstein

**Affiliations:** Dept. of Biomedical Engineering, University of Michigan, Ann Arbor, USA; Dept. of Neurosurgery, University of Michigan Medical School, Ann Arbor, USA; Microscopy Core, Biomedical Research Core Facilities, University of Michigan, Ann Arbor, USA; Laboratoire PCC, UMR168, Institut Curie, PSL, CNRS, Sorbonne Université, 75248 Paris, France; Department of Neurosurgery, NYU Langone Health, New York, NY, USA; Dept. of Cell and Developmental Biology, University of Michigan, Ann Arbor, USA; Dept. of Physics and Astronomy, Johns Hopkins University, Baltimore, USA

## Abstract

Active nematic liquid crystals are the main structural phase of gliomas, promoting collective migration and aggression. We establish the existence of nematic order and topological defect lines and loops in 3D in vivo mouse and human glioma brain tumors. As predicted by theory, sections through the disclination lines in 3D appear as ±1/2 topological defects in 2D. In 3D, these defects either persist along disclination lines or twist as they interconvert from −1/2 to +1/2. Cell alignment exhibits quasi–long-range order, spreading throughout the tumor over distances between 300−3000 *μ*m. In vitro −1/2 and +1/2 defects display changes in apoptosis levels, suggesting topological defects regulate glioma cell density. The large scale order of gliomas correlates with tumors’ aggressive behavior. The organization of gliomas as active nematic liquid crystals provides a novel physical foundation of complex solid tumors; their deconstruction signposts potential treatments for deadly cancers.

---

Whether gliomas consist of random accumulations of cells or are self-organizing remains unknown. If large scale order exists, it should manifest as invariant structures across different tumors. Active liquid crystals form a class of soft materials in which the underlying units have anisotropic shapes, typically rod-like, and form highly aligned configurations, yet also exhibit singular configurations like disclinations (*1, 2, 3, 4*). As active matter, the unit cells take in energy and convert it to forces, resulting in bulk, collective motion and emergent behavior. In recent works, it has been found that certain cell types in 2D in vitro, including epithelial, neuroepithelial, fibroblast, myoblasts and outer epithelium of *Hydra*, display nematic alignment and topological defects, characteristic of crystalline order (*5, 6, 7, 8, 9, 10, 11*). It has been proposed that crystalline order influences tissue morphogenesis, cellular extrusion and apoptosis, and mechanotransduction (*5,11,12,13,14,15,16*). These results have been obtained in 2D geometries and 2D surfaces that can deform in 3D. Work on the invasive front of breast cancer, mostly in 2D, recently suggested that nematic alignment is present within cancer cells and topological defects are present within the surrounding extracellular matrix (ECM) (*17*). Whether topological structures exist within tumor bulk in a 3D in vivo pathophysiological context remains to be established.

In this work, we show that glioma brain tumors in vivo, and in vitro, are structured as active nematic liquid crystals. Building on our previous work that gliomas exhibit self-organized, aligned, multicellular structures, termed oncostreams (*18*), we demonstrate that gliomas display nematic order, topological defects, disclinations, and quasi-long range order in 2D and in 3D. Significantly, the amount of nematic order scales with tumor aggression - suggesting crystalline order contributes to tumor malignancy - constituting a novel potential therapeutic target for this incurable cancer.

## From small- to large-scale nematic spatial organization in gliomas

Gliomas display hallmarks of nematic phases: mesoscopic nematic alignment, topological defects, and quasi-long range order (QLRO). To study the large-scale 3D spatial organization in brain tumors in vivo, we reconstructed 3D sequential sections of hematoxylin and eosin (H&E) stained mouse and human tumors (Fig. 1A) and optically-cleared mouse tumors imaged directly in 3D with light-sheet microscopy (LSM) (Fig. 1B). In 2D geometries, like an H&E section of glioma-bearing mouse brains or an optical plane of cleared glioma-bearing mouse brains, mesoscopic nematic alignment was previously found in oncostreams as cells aligned along a common orientation with back-and-forth motion ((*18*), and Fig. 1, C and E). In nematic liquid crystals, the local direction of alignment is described by the director field (*19*) and topological defects are singular configurations of the director field with a non-vanishing winding number. We detected numerous examples of such defects in 2D geometries as comets (+1/2 winding number) or trefoils (−1/2 winding number) (*19*). Cellular arrangements that resemble trefoil defects were mainly found at the intersection between two or more oncostreams (Fig. 1, D and G).

**Figure 1:**
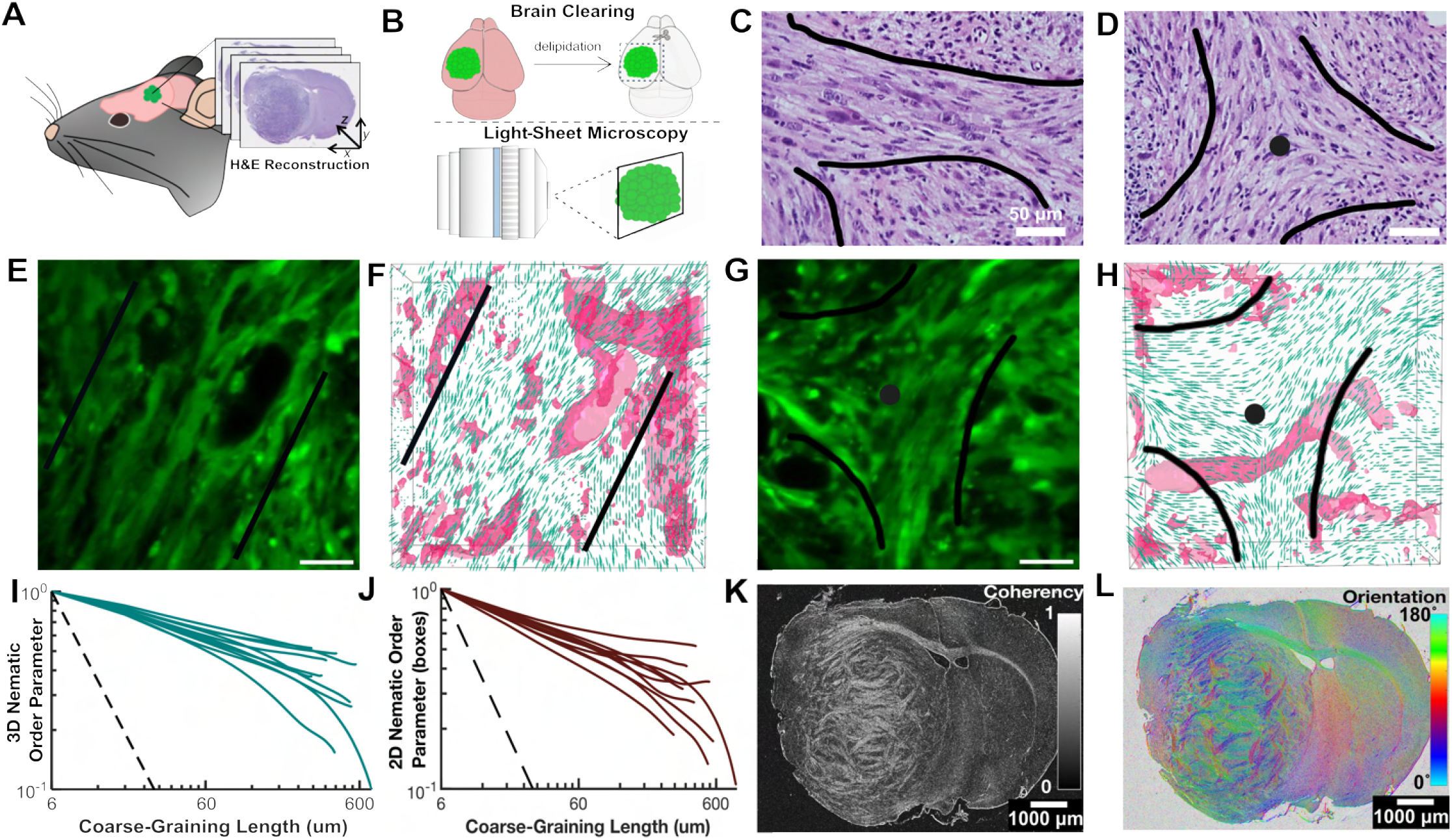
2D and 3D nematic order and topological defects in whole mouse gliomas. (**A**) Schematic of hematoxylin and eosin (H&E) 3D reconstructions of mouse brain and tumor from microtome sectioned serial 5 µm slices of paraffin embedded tissue (see Materials and Methods for more details). (**B**) Schematic of brain clearing, delipidation, tumor dissection, and 3D light sheet microscopy (LSM) imaging of GFP-tagged cleared fixed whole brains from glioma-bearing mice (see Materials and Methods for more details). (**C**) H&E stain of a region of high nematic order (oncostream) shown in 2D (inside black lines) from the mouse glioma reconstructed in panel A. (**D**) H&E stained section of the mouse glioma in panel A containing a topological defect (black lines roughly outline cell orientation and singularity marked with a dot). (**E**) Oncostream region (inside black lines) imaged with LSM from a cleared fixed whole mouse glioma. (**F**) 3D director field (green ellipsoids) corresponding to panel E. Blood vessels are outlined as pink surfaces. (**G**) Topological defect (black lines outline cell orientation and singularity marked with a dot) imaged with LSM. (**H**) 3D director field (green ellipsoids) corresponding to panel G. Blood vessels are outlined as pink surfaces. (**I**) 3D nematic order parameter against coarse-graining length for the NPD tumor sample in panels E-H. Each teal curve refers to a different ROI from the sample. (**J**) 2D nematic order parameter against coarse-graining length for the NPD tumor sample in panels E-H. Each red curve refers to a different ROI from the sample. (**K**) Coherency map of a full mouse brain and tumor H&E section (see Materials and Methods for the description of coherency). Color bar corresponds to the magnitude of the coherency. (**L**) Orientation angle color map of brain in (K). Color bar corresponds to the orientation angle. Scale bars: C, D, E, F – 50 *μ*m. K, L – 1000 *μ*m.

To integrate the third dimension of gliomas, we determined the director field from high-resolution 3D imaging data of high-grade mouse gliomas (Fig. 1B, see Materials and Methods). Two cross-sections of the director field (green ellipsoids) are shown in Fig. 1, F and H, corresponding to the imaging planes in Fig. 1, E and G, respectively. To assess the degree of nematic order, we computed the 3D nematic order parameter, *S*_3*d*_, in cubic domains of varying size (coarse-graining length). This order parameter is zero for a purely random configuration and one for a perfectly aligned configuration (see Materials and Methods). We found that *S*_3*d*_ follows a power law decay, which is characteristic of QLRO, over domain sizes above 300 *μ*m (Fig. 1I). The fitted exponent was −0.29 ± 0.06 (mean±std, *n* = 10). This shows that the large-scale spatial organization of gliomas features 3D nematic alignment.

A projection of the 3D nematic alignment onto a plane illustrates both 2D nematic alignment, oncostreams (*18*), (Fig. 1, C and E), and configurations that resemble +1/2 and −1/2 topological defects (Fig. 1, D and G). Furthermore, the 2D nematic order parameter, *S*_2*d*_, qualitatively reproduced the behavior of *S*_3*d*_ as a function of the coarse-graining length (Fig. 1J). Importantly, the fitted parameters of the power law were comparable (−0.36 ± 0.1 vs. −0.29 ± 0.06 (*n* = 10)). This suggests that the 2D nematic order faithfully represents the 3D nematic alignment.

Moving to larger lengthscales on the order of millimeters, we found that tumor regions display a significantly larger nematic order than in normal surrounding brain regions (i.e., striatum and neocortex) in H&E samples (Fig. 1K). Although nematic order varied between tumor areas, large areas of absent order were not encountered, suggesting that tumors are mostly ordered. Interestingly, the *corpus callosum*, the bundle of axons connecting both hemispheres, displayed a high nematic order, likely the result of the parallel orientation of the axon bundles and oligodendrocytes. The further existence of crystalline order and topological defects in the *corpus callosum* was not studied further. Additionally, in tumor regions we found nematic domains of sizes from hundreds of microns to millimeters (Fig. 1L).

### Quasi-long-range nematic order in gliomas

To further characterize the large-scale nematic organization of gliomas, we analyzed the nematic order parameter for various sub-types of mouse and human tumors and brain regions (see Materials and Methods). From 3D reconstructions of sequential H&E sections, various tumor regions of interest (ROI) and normal brain ROI were analyzed. A table of all brains imaged and full H&E images are shown in fig. S1E. The high-grade mouse glioma models -NPD and NPA - and human gliosarcomas are known to be more aggressive compared to the low-grade mouse glioma model, NPAI (tumor model genetic specifications found in Materials and Methods, and survival curves shown in fig. S2) (*18, 20, 21, 22*). Additionally, we analyzed optically-cleared NPD mouse tumors imaged with LSM to obtain high resolution in all three dimensions (see Materials and Methods, Movie S1).

Due to the asymmetric resolution of sequential H&E sections (sub-micrometric in *x* and *y* and 5 − 6 *μ*m in *z*), we utilize the 2D nematic order parameter *S*_2*d*_ as a proxy for determining the 3D nematic organization (see Materials and Methods). Here, we tested two averaging domains: squares (Fig. 2A, C, and E) or boxes (Fig. 2B, D, and F). Both domains were defined by a single coarse-graining length *ℓ* (Fig. 2, A and B). Note that in squares, the in-plane director field alignment within a single image plane is evaluated, whereas in boxes, alignments from surrounding image planes are also incorporated.

**Figure 2:**
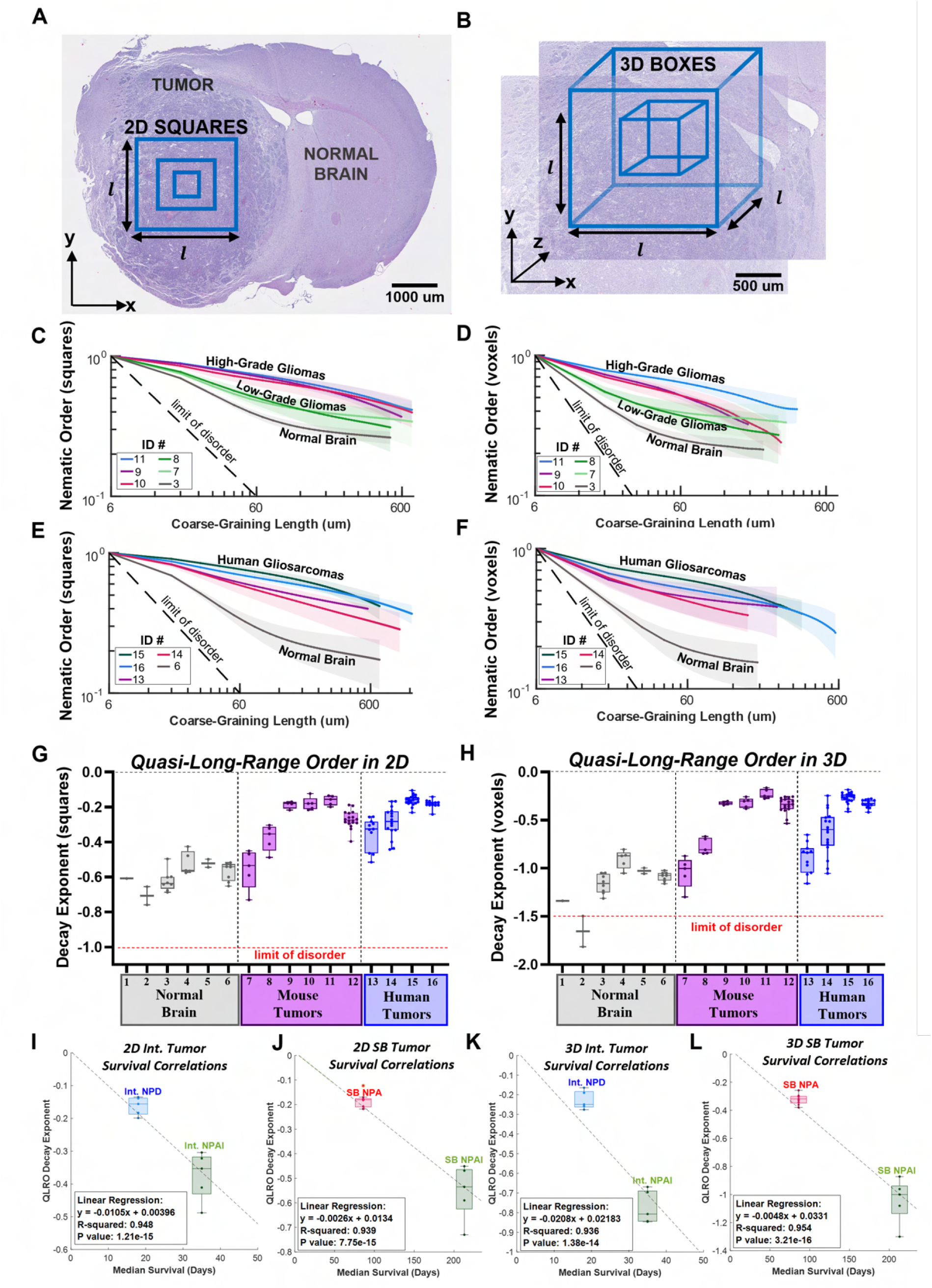
2D and 3D quasi-long range order (QLRO) in in vivo mouse and human gliomas. (**A**) An example H&E section of mouse glioma NPD tumor. The blue 2D squares represent regions of interest (ROI) for the computation of the 2D nematic order parameter (NOP) (see Materials and Methods for definition). The parameter *l* represents the coarse-graining length and corresponds to increasing sizes of the squares, scale bar = 1000 µm. (**B**) An example ROI in the H&E section of the NPD tumor in A. The blue 3D boxes represent regions for the computation of 3D NOP, scale bar = 500 µm. (**C-D**) Respectively, 2D and 3D NOPs (see Materials and Methods for definitions) as a function of the coarse-graining length (see panels A, B). The dashed line indicates the theoretical limit of a random disordered case. Colored curves show NOP for various mouse gliomas, as indicated in the legend (ID numbers found in bottom table of fig. S1). Red, blue, and purple curves are high-grade gliomas; light and dark green curves are low-grade gliomas; and gray curve is normal brain. (**E-F**) Respectively, 2D and 3D NOPs (see Materials and Methods for definitions) as a function of the coarse-graining length. The dashed line is the theoretical limit of a random case. Red, blue, green, and purple curves show NOP for various human gliosarcomas, indicated in the legend, and the gray curve shows NOP for tumor-adjacent normal brain regions. (**G-H**) Respectively, 2D and 3D decay exponents for various tumor and normal brain ROI. For each ROI, the NOP as a function of the coarse-graining length was fitted with a power law: *f* (*x*) = (1 − *c*)*x*^*b*^ + *c* (see Materials and Methods for details). The decay exponent corresponds to the fitting parameter *b*. Box and whisker plot shows each ROI’s exponent with mean shown as horizontal line. X-axis labels correspond to each tissue listed in the table in fig. S1. The horizontal red lines correspond to the exponent in the theoretical limit of a random case. (**I-L**) Respectively, correlations of 2D and 3D QLRO with mouse median survival for Intracranial and Sleeping Beauty mouse models. Abbreviations for tumor ID names defined in Materials and Methods. All brain samples and corresponding ROI shown in fig. S1. In all panels, shaded areas correspond to the standard deviation over multiple ROIs.

For a fixed coarse-graining length, the nematic order parameter *S*_2*d*_ in low-grade and high-grade mouse gliomas and human gliosarcomas is larger than the control normal brain (Fig. 2, C-F). Additionally, the tumors’ order again propagated significantly farther than in tumor-adjacent (T-A) normal gray matter brain regions of each mouse and human samples, as indicated by the higher levels of nematic order in the tumors (figs. S3-S6). Furthermore, mouse high-grade gliomas predominantly exhibited larger magnitudes of the nematic order compared to low-grade gliomas (Fig. 2, C and D).

The nematic order parameter decays monotonically with the coarse-graining length (Fig. 2, C-F). This is expected as its magnitude tends to decrease as the length of the averaging domain increases due to, for instance, de-correlation of nematic alignment or topological defects (*19, 23*). Importantly, all cases deviate from the theoretical limit of a random case, *S* ∼ *ℓ*^−*d*/2^, where *d* is the dimensionality of the averaging region (*d* = 2 squares, and *d* = 3 boxes), see Methods, confirming nematic correlations.

To further characterize the spatial decay of nematic alignment, we fitted the nematic order parameters with a power law with a constant deviation term, *f* (*ℓ*) = (1 − *c*) * (*ℓ*/*ℓ*_*c*_)^*b*^ + *c*, see Methods. The minimal cut-off length *ℓ*_*c*_ = 6 *μ*m was fixed. This function describes two limit cases that are controlled by a crossover length *ℓ*_*m*_ ∼ *ℓ*_*c*_ (*c*/(1 − *c*))^1/*b*^: for sufficiently small coarse-graining length *ℓ* ≪ *ℓ*_*m*_, *f* (*ℓ*) decays as a power law with a negative decay exponent *b* < 0, which is characteristic of quasi-long range order. The decay exponent is expected to be non-universal and to depend on sample properties (*24, 25*). For sufficiently large coarse-graining length *ℓ* ≫ *ℓ*_*m*_, the nematic order parameter is independent of the coarse-graining length, which is characteristic of long-range order. The parameter *c* quantifies the asymptotic level of order at large length scales. In the following we refer to quasi-long range order as the cases where the nematic order parameter decays with the distance in a manner compatible with a power law. Alternatively, the cases where the nematic order parameter saturates asymptotically to a constant value are referred as long-ranged order (LRO).

The fitted values of all decay exponents are summarized in Fig. 2, G and H. We found that the decay exponent is smaller (less negative) for the human gliosarcomas, mouse NPA, and mouse NPD tumors compared to mouse NPAI (*P* value < 0.0001). Further, all tumor cases have significantly smaller decay exponents compared to the normal gray matter brain regions (NPA, NPD, gliosarcoma: *P* value < 0.0001; NPAI: *P* value = 0.007, all compared to normal brain). The tumor-adjacent normal gray matter brain regions have significantly higher (more negative) decay exponents than their neighboring tumors (Fig. 2, G and H, and figs. S3-S6).

As NPAI tumors are significantly less aggressive than NPA and NPD tumors (fig. S2, (*26*)), we wanted to establish whether median survival correlates with the decay exponent. Using a linear regression model, we found a negative correlation between median survival and the decay exponent, indicating that a smaller decay exponent, and thus, a higher propagation of nematic order, confers increased tumor aggression (Fig. 2, I-L). These data correlate with our previous results showing that higher density of oncostreams, the locus of nematic order in gliomas, correlates with tumor malignant behavior (*18*).

The fitting procedure also provides an estimate of the parameter *c*, which varies from zero (no order) to one (perfectly ordered). Whereas *b* describes the existence of QLRO, *c* describes the remaining, much lower, long-range nematic order. In the control normal brain gray matter, the value of *c* was 0.2 ± 0.04 (mean±std, n=8), and tumor-adjacent normal brain regions *c* ranged from 0.2 −0.5 (fig. S7). This suggests that the normal brain has a modest intrinsic nematic order. Notably, we found that in low-grade gliomas, the value of *c* was similar to tumor-adjacent brain regions, while it was nearly vanishing for most of the high-grade gliomas analyzed (fig. S7). Altogether, this data shows that the spatial propagation of nematic order in high-grade gliomas obeys a power law decay for a length scale that ranges up to a few millimeters. In conclusion, nematic order in high-grade gliomas is quasi-long range, contrasting with the low levels of long-range order exhibited in the normal brain.

### Gliomas exhibit disclination lines and loops

Another property of nematic phases in 3D geometries is the existence of disclinations - line-like singularities in the director field, around which the director rotates by 180 degrees in a plane normal to a specific direction called the rotation vector **Ω** (*23, 27*). To study disclinations in whole gliomas in 3D, we focused on optically-cleared mouse tumors imaged with LSM and visualized with open-ViewMin (*28*) (Fig. 1B, Movie S1).

Surfaces bounding regions of low nematic order (purple domains in Fig. 3) have a serpentine tubular geometry. These surfaces can merge with other surfaces, split, form closed loops, or connect interfaces of vessels (pink domains in Fig. 3 and figs. S8-S10). As a result, they form an intertwined network throughout the tumor environment.

**Figure 3:**
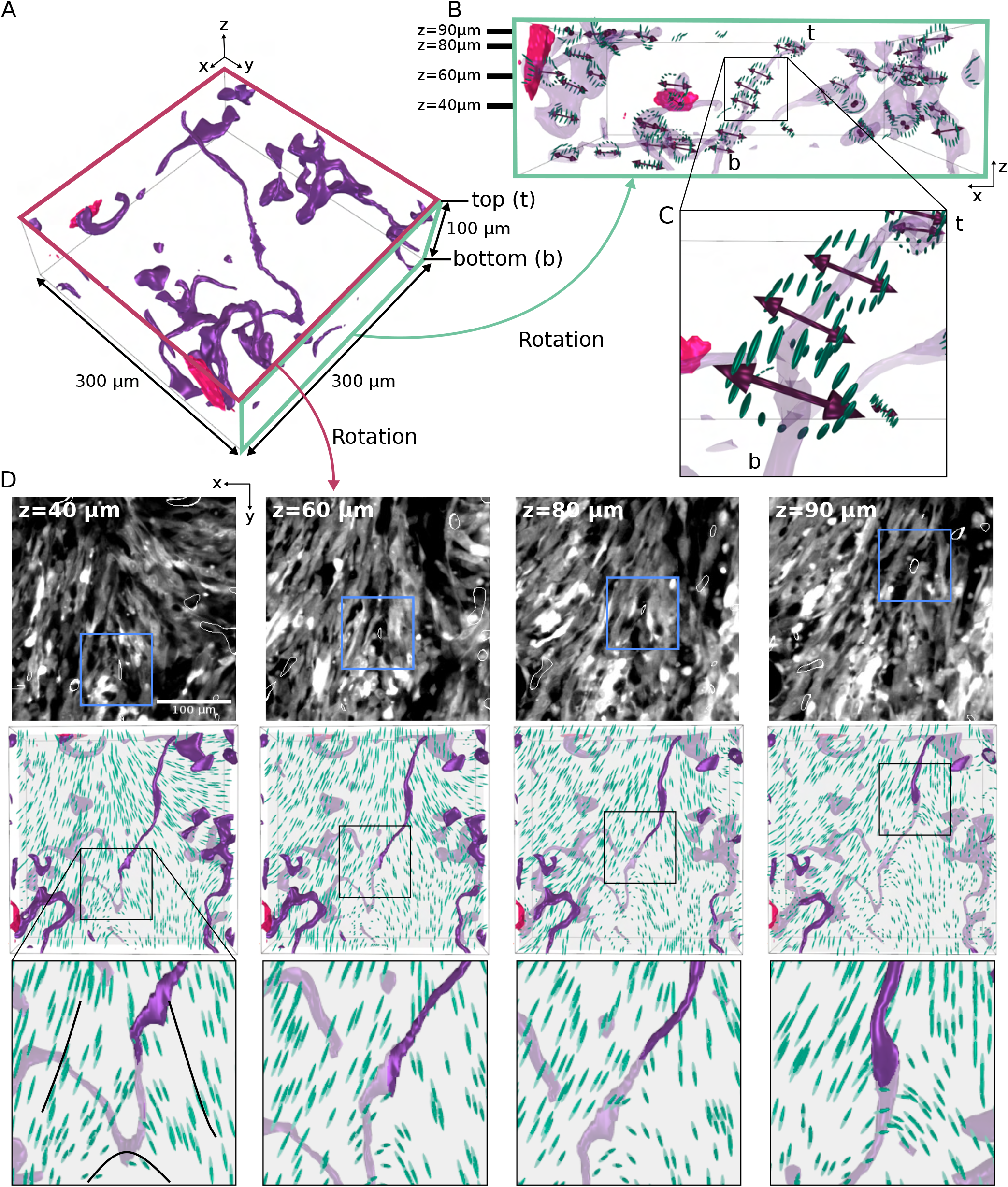
3D topological defects in in vivo mouse gliomas. **(A)** 3D reconstruction of a region (300 × 300 × 100 *μ*m^3^) from cleared intracranial NPD mouse glioma. Dark purple regions correspond to surfaces with low nematic order (see methods). Pink regions outline blood vessels. The top *z*-plane is labeled “t” and the bottom plane is labeled “b.” **(B)** Side view of the 3D reconstruction in panel A. The green rectangle corresponds to the green face in panel A. To facilitate visualization, the dark purple areas in panel A are now translucent purple. The 3D director field encircling the purple regions is shown as green ellipsoids and the double-headed red arrows represent the rotation vector, Ω. The short black lines on the top left corner of the green square label the *z*-planes that are shown in D. (**C**) Approximate zoom in of the black square in panel B. (**D**) Top row: Gray-scale LSM images of cleared tumor with blue boxes outlining disclination lines (thin white outline) through the *z*-plane (*z* coordinate indicated in top left corner). Middle row: Corresponding 3D director field (green ellipsoids) reconstructions with purple regions corresponding to surfaces with low nematic order and disclination-line locations now outlined with black boxes. Note: the *z*-plane is translucent gray, and thus the features above the *z*-plane appear in a darker color and the features below the *z*-plane in a dimmer color. Bottom row: Zoomed in view of middle row boxes showing disclination lines traveling through the *z*-plane. The scale bar is 100 *μ*m.

The configuration of the director field around low-nematic order surfaces reveals the existence of disclinations (Fig. 3, B and C). In some cases, the rotation vector **Ω** (double-headed arrows in Fig. 3, B and C) is nearly perpendicular to the main direction of the low-nematic order surfaces, indicating a twist profile (Movie S2). Remarkably, twist-like disclinations confirm the 3D nature of topological defects and demonstrate the necessity of the 3D nematic alignment. When scanning across *z*-planes (Fig. 3D, top panels), disclinations can be located at the intersections between two or more oncostreams, forming a trefoil-like director field configuration (Fig. 3D, middle and bottom panels). Here, the director field maintains a trefoil-like configuration along multiple z-sections of the disclination (Fig. 3D, Movies S1 and S3). However, due to the twist-like nature of this disclination, the director field configuration can resemble a comet-like topological defect, when the same disclination is viewed from another angle (fig. S10, Movie S4), further emphasizing its 3D nematic nature.

In other disclinations (figs. S8), the director configuration changed from a trefoil to a comet along the disclination line, for instance when the disclination bifurcates or forms a loop. Finally, we also observed wedge-like disclinations where the rotation vector **Ω** is nearly parallel to the main direction of the low-nematic order surfaces (fig. S9)

To further complement the above analyses, we turned to 3D reconstructions from sequential H&E sections and identified regions in high-grade tumors that appear like trefoils or comets. Numerous examples of trefoil configurations were found in NPA and NPD mouse models (figs. S11-S14). These arrangements propagated across several image planes with lengths ranging from 15 to 60 *μ*m (figs. S11-S14). With less frequency, comet-like cellular arrangements were also found in NPD mouse models with lengths between 30 to 65 *μ*m (fig. S15). Such patterns were not found within normal brain and tumor-adjacent brain sections. Altogether, our data shows that 3D disclinations are emergent structures across multiple tumor types.

### Topological defects influence cell density in gliomas

Past works have reported that several biological processes including cell extrusion, cell accumulation, and cell death can occur preferentially near topological defects (*29, 30*). To evaluate the potential function of topological defects, we utilized an in vitro glioma cell culture platform (fig. S16) (*31*). In this 2D platform, both −1/2 and +1/2 topological defects formed over 36 − 48 hours post-seeding (Movies S5-S6). Time course analysis showed that the overall nematic order parameter *S* tended to increase, while the density of topological defects tended to decrease (fig. S16). Additionally, we found that topological defect pairs of +1/2 and −1/2 can annihilate each other (fig. S16G) or remain stable (fig. S16H). Since many topological defects remain, the cell culture never reaches 100% nematic alignment (*S*_2*d*_ = 1). Finally, +1/2 defect motion was predominantly tail-to-head (fig. S17).

To assess the role of topological defects in cell death, we utilized a fluorescence apoptosis marker, NucView 530 Caspase-3, in our time-lapse movies to obtain a spatiotemporal view of apoptosis (Movies S7, S8). To determine the distribution of apoptosis, we compared the apoptosis fluorescent values in randomly placed square ROIs to the same size square ROIs around topological defects (Fig. 4A). Apoptosis was drastically increased at −1/2 topological defects, by about 30%, and decreased at the head of +1/2 comets by about 33% (Fig. 4, B and D). Additionally, we determined cellular density using the nuclear dye Hoescht. −1/2 topological defects displayed an 11% lower cell density at the defect core (Fig. 4, C and E). Interestingly, cell density is also decreased by about 7% at the +1/2 defects, indicating that apoptosis and cellular density are not always inversely related and other mechanisms may decrease cell density, such as collective cell migration.

**Figure 4:**
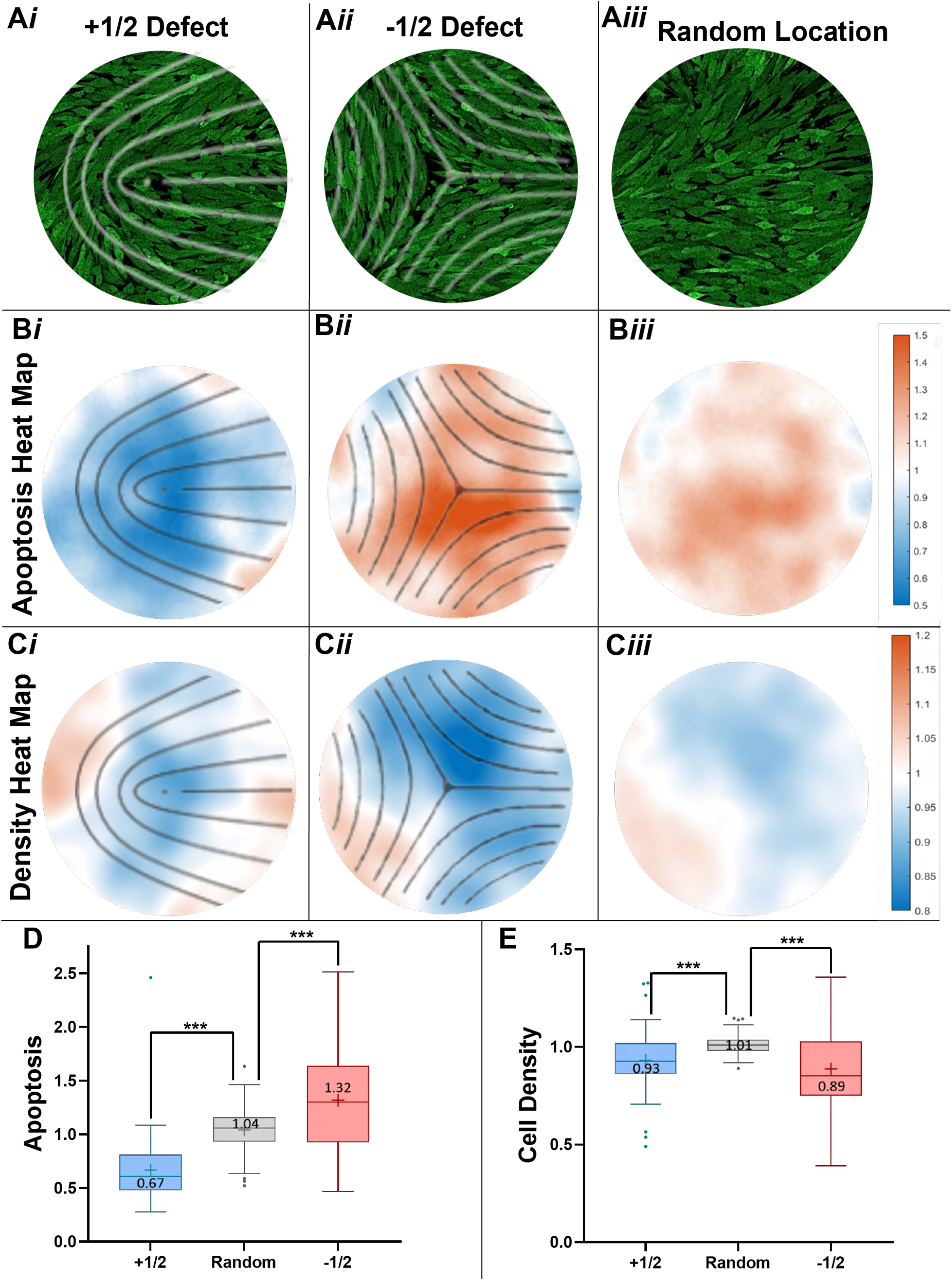
Function of topological defects in 2D glioma cells in vitro. (**A**) Confocal images of in vitro glioma cell cultures that display different nematic organization patterns: (i) +½ topological defects (overlaid in white lines), (ii) -½ topological defects (overlaid in white lines), and (iii) random location within the culture. The diameter of the circles is 224 µm. (**B**) Heat maps of average fluorescent signal of Caspase-3, indicating apoptotic cells (see Materials and Methods) for the three regions in panel A. The signals are normalized by the average of the entire field-of-view and defects are now overlaid with black lines. (**C**) Heat maps of average number of cell nuclei (stained with Hoechst, see Materials and Methods), indicating cell density for the three types of regions in panel A. The signals are normalized by the average of the entire field-of-view. **(D)** For each subpanel of B, the normalized fluorescence intensity inside a centered square region of side length 116 µm was averaged and plotted in a box and whisker plot of apoptosis (means labeled as +). (**E**) For each subpanel of C, the normalized number of cell nuclei inside a center square region of side length 116 µm was averaged and plotted in a box and whisker plot of cell density (means labeled as +). Panels D and E: n=52 for +½ defect, 45 for -½ defect, and 97 for random location with N = 3 movies. *P* values *** < 0.005.

In this study, we investigated the high-order 3D organization of glioma tumors, finding that gliomas display nematic order, topological disclinations, and quasi-long range order. Additionally, increased order is positively correlated with tumor aggression. Thus, the aforementioned crystalline properties of gliomas are possible viable therapeutic targets to treat one of the most aggressive and devastating cancers with a two-year survival rate of less than 5%, as clinical strategies have remained stagnant since 2005 (*21, 22, 32*). Targeting the global physical order of glioma organization is an exciting and novel therapeutic approach.

## Acknowledgments

Research reported in this publication was supported by the National Cancer Institutes of Health under Award Number P30CA046592 by the use of the following Rogel Cancer Center Shared Resource: Tissue and Molecular Pathology Shared Resource Molecular Pathology Research Laboratory. Additionally, this project has received funding from the European Union’s Horizon 2020 research and innovation programme under the Marie Sklodowska-Curie grant agreement No 847718. This publication reflects only the author’s view and that the European Research Agency is not responsible for any use that may be made of the information it contains. We thank Gabriel Corfas, Jacques Prost, Robin Selinger, Jonathan Selinger, and Suraj Shankar for comments on the manuscript.

## Funding

National Science Foundation Graduate Research Fellowship Grant DGE-1841052 (AEA). University of Michigan Rackham Predoctoral Fellowship (AEA). National Institute of Biomedical Imaging and Bioengineering (NIH/NIBI): R01-EB022563 (PRL, MGC). National Cancer Institute (NIH/NCI) U01CA224160 (MGC). Rogel Cancer Center at The University of Michigan G023089 (MGC). Ian’s Friends Foundation grant G024230 (PRL). Leah’s Happy Hearts Foundation grant G013908 (PRL). Pediatric Brain Tumor Foundation grant G023387 (PRL). Chad-Tough Foundation grant G023419 (PRL). NSF CAREER Award DMR-2225543 (DAB). Marie Sklodowska-Curie Grant No 847718 (QC). National Cancer Institutes of Health Award Number P30CA046592 (Rogel Cancer Center Shared Resource: Tissue and Molecular Pathology Research Laboratory).

## Author contributions

AEA, CBM, and PRL conceived the research. PRL, CBM, and MGC supervised the project. AEA, PRL, MLV, GS, MER, MBE, DPV, CBM designed and performed the experiments. GS and DPV generated mouse tumors and performed all animal work. BJ instructed and performed brain clearing and light-sheet microscopy. DAO and JAH provided the human patient gliosarcoma samples. AEA, QC, DAB, CBM, and PRL analyzed the data. AEA, CBM, and PRL wrote the manuscript (with input from all coauthors).

## Competing interests

There are no competing interests to declare.

## Data and materials availability

Experimental data and the code to detect and analyze the 2D topological defects are available on Dryad (10.5281/zenodo.15199294). The code to compute the director field in 3D is available in (*13*).

## Supplementary materials

Materials and Methods

Figs. S1 to S17

Movie S1 to S8

## Supplementary Materials for

### Materials and Methods

#### In Vivo Mouse Glioma Generation

##### Genetically Engineered Mouse Glioma Models (GEMM)

All in vivo experiments were conducted according to the guidelines approved by the Institutional Animal Care and Use Committee (IACUC) at the University of Michigan protocols PRO00011290, PRO00011291, and PRO00011292. All animals were housed in an AAALAC accredited animal facility at the University of Michigan. Animals were monitored daily. All tumor bearing animals were euthanized at the time they develop clinical signs of tumor burden. Maximal tumor burden did not exceed IACUC guidelines. Tumors harboring different genetic drivers were generated with the Sleeping Beauty (SB) transposon system, following our methodology (*20, 26*). Genetic models included the following gene expressions and inhibitions: (i) *shP5*3, *NRAS-G12V*, and *shAtrx* (NPA), *(ii) shP53, NRAS-G12V*, and *Pdgfβ* (NPD), (iii) *shP5*3, *NRAS-G12V, shAtrx*, and *IDH1-R123H* (NPAI), as in (*18, 20, 26*). Tumors were also tagged with endogenous green-fluorescent protein (GFP) for visualization. Tumors are labeled as “SB” in fig. S1E.

##### Intracranial Implantable Syngeneic Mouse Gliomas

Intracranial glioma tumors were generated by stereotactic intracranial implantation into the mouse striatum of 3.0×10^4^ mouse glioma neurosphere cells (NPA, NPD, NPAI) in 6–8 week old females C57BL/6 mice (Taconic Biosciences) following previous methodology (*18, 20, 26*). Tumor cells are also tagged with endogenous GFP for visualization. Tumors are labeled as “Int.” in fig. S1E.

#### H&E tumor sections

Mice were transcardially perfused with oxygenated Tyrode’s solution, followed by 4% buffered paraformaldehyde and the brains paraffin-embedded. Brains were microtome sectioned in 5 *μ*m thick serial sections and then stained with hematoxylin and eosin (H&E) to visualize extracellular matrix and cell cytoplasm and nuclei.

#### Brightfield Whole-Slide Imaging

Brightfield whole-slide scanning of serial H&E sections was performed by the University of Michigan Tissue and Molecular Pathology Shared Resource (TMPSR) Molecular Pathology Research Laboratory (MPRL) on the Vectra Polaris at 40x magnification.

#### Human Patient Samples

FFPE human gliosarcoma (GSC) samples were obtained from primary surgery from the University of Michigan Medical School Hospital. Patients gave informed consent for collection of tissue collection under Institutional Review Board-approved Protocol (HUM00057130) at the University of Michigan. All patients were part of the clinical trial detailed in (*33*). Samples referred to as GSC #1 and #2 are from Patient #19, GSC #3 is from Patient #20, and GSC #4 is from Patient #12 in the clinical trial. Samples were sectioned, H&E stained, and imaged as described above.

#### 3D In Vivo Light Sheet Scanning Microscopy

##### Brain Clearing and Immunolabeling

To clear and label brains, the Life Canvas Active Clarity protocol was followed (*34, 35*). Mice were terminally anesthetized and perfused with oxygenated Tyrode’s Solution, followed by hydrogel monomer (HM) solution containing 4% acrylamide, 0.05% bisacrylamide, 4% paraformaldehyde (PFA), 0.25% VA-044 thermal initiator in phosphate buffered saline (PBS). Brains were extracted, halved, and sectioned around the tumor, then incubated for 3-7 days at 4°C in HM solution. The brains were then purged with nitrogen gas under vacuum, then incubated overnight at 37°C. Once the solution was gelled, brains were rubbed out of the gel and incubated overnight in delipidation buffer (Life Canvas Buffer A) at 37°C. To clear lipids, brains were then placed in mesh bags, then into the Life Canvas machine with Buffer A and run for 72 hours with electrophoresis at 1000 mA. After clearing, the samples are washed with PBS at RT. To boost endogenous green fluorescence protein (GFP) signal of the glioma cells within the brain, immuno-labeling with rabbit Anti-GFP antibody (Abcam ab290) was necessary. Briefly, samples were incubated in primary sample buffer overnight, then stained for 16 hours with primary antibody and 1000 mA electrophoresis. After PBS washes, samples were fixed with 4% PFA overnight. Samples were then incubated in secondary sample buffer and then stained for 8 hours with secondary antibody, Alexa Fluor Donkey Anti-Rabbit 488 (Fisher Scientific PIA32790), washed with secondary sample buffer alone, then PBS, and fixed overnight in 4% PFA. Index matching was performed for 24 hours in 50% EasyIndex (refractive index = 1.52) in PBS, then for 2-3 days in 100% EasyIndex. 1 µg/mL DAPI was added in the EasyIndex for nuclear staining. After refractive index matching, brains appeared translucent and barely visible within the solution.

##### Light Sheet Microscopy

A Zeiss Light Sheet 7 with a 20x immersion objective lens was utilized to perform broad-range *z*-stack light sheet microscopy at the University of Michigan Biomedical Research Core Microscopy Facility. The light sheet was aligned on both the right and left sides of the microscope. Large sections of tissue were imaged throughout tumor pieces approximately 1 cm^3^ in size.

#### In Vitro 2D Experiments

##### Glioma Cell Lines and Culture Conditions

Mouse glioma cells were maintained at 37°C with 5% CO_2_ and DMEM/F-12 media with Normocin, N2, and B27. Neurospheres were derived from our GEMMs. To produce adherent cells for in-vitro imaging, NPA neurospheres were cultured on laminin coated flasks with DMEM + 10% fetal bovine serum and used after three passages.

##### In Vitro Oncostream Platform

For the in vitro platform, glass coverslips or 35 mm glass-bottom culture dishes (Ibidi) were coated with Poly-D-lysine and laminin proteins as in (*31*). Culture surfaces were first acid washed with 1M hydrochloric acid for 2-4 hours at 60°C, then rinsed with milli-Q-water, and 100% ethanol. This removed dust or contamination that could possibly create artificial cell patterns. Then, poly-D-lysine at a working concentration of 100 µg/ml was applied to the culture surfaces overnight at 4°C. After drying and rinsing with sterile PBS, laminin at 100 µg/ml diluted in PBS with MgCl and CaCl was applied and incubated for 2 hours at 37°C, then at 4°C to solidify overnight. Before cell seeding, the dish was rinsed twice with cold PBS with MgCl and CaCl and once with PBS without MgCl and CaCl. Adherent NPA cells were seeded at a density of 2×10^5^ for a 35 mm dish and incubated for 24 hours at 37°C to allow for adherence.

##### Apoptosis Cell Labeling

To track cellular apoptosis in time-course movies, we utilized NucView 530 Caspase3 (Biotium 10406-T) according to the manufacturer’s protocols. Time lapse imaging was performed as normal with the addition of the red wavelength laser.

##### Time-Lapse Imaging

For time lapse imaging, we utilized an inverted Zeiss LSM880 laser scanning confocal microscope with AiryScan (Carl Zeiss, Jena, Germany) equiped with an incubation chamber set to 37°C and 5% CO_2_. Cells were imaged periodically (with time interval ranging between 10-46 min), for 200-360 cycles (26 - 88 h) at 3×3 tiles at 10x or 20x resolution. We imaged 6 cultures, imaging 6-9 locations each, (n=6 movies, N=40 imaged positions).

##### Post-Processing of Time-Lapse Confocal Movies

After imaging cultures for up to 3 days, we utilized the FIJI plug-in OrientationJ to create vector fields of orientation for each image in the time-lapse movies. Before processing, we binned each image 2×2×1 to optimize image data size. For images taken with the 20x lens, we utilized a local window sigma of 30 px, Gaussian gradient, and 30 px grid size (25 *μ*m). For images taken with the 10x lens, we utilized a local window sigma of 15 px, Gaussian gradient, and 15 px grid size. We then calculated the local order parameter, *Q*_*loc*_ over square sub-windows of size sigma=8 um (Eqn. 2). As there are points where alignment is disrupted, *Q*_*loc*_ = 0 at topological defects while *Q*_*loc*_ = 1 at perfectly aligned regions. We then created a local order parameter heatmap to visualize where *Q*_*loc*_ = 0. We found that *Q*_*loc*_ was about 1 everywhere in the images except for small, confined areas where *Q*_*loc*_ tended to 0 at the center of a topological defect. When overlaying the heatmaps onto the original images with vector fields, we found the defect locations which were corroborated visually. For the apoptosis localization analysis, the Caspase-3 channels were thresholded 24:255 on ImageJ, then Gaussian Blur 15 px was used. For spatial density analysis, the command “Make Binary” was used, and then “Gaussian Blur” set to 15 px.

#### Alignment of Sequential H&E Sections

Sequential H&E sections were aligned by either of the two following procedures:

1. For a given H&E section, the next H&E section was transformed by a combination of translations and rotations until optimal matching between the two sections was achieved. The criteria for optimal matching was visual inspection. This procedure was repeated until the whole series of H&E sections was aligned.
2. We used the script that is described in Ref. (*36*) to align sequential H&E sections in an automatic way. The script is available on the following GitHub page: https://github.com/ashleylk/CODAI. In short, this script performs a combination of transformations such as translations, rotations or interpolations to match pairs of consecutive H&E sections. Finally, the alignment of the output series of H&E sections was further improved by applying Procedure 1 on them.

The brain tumors (see fig. S1E for ID numbers): SB NPAI #1 (ID 1 and 7), Int. NPAI #2 (2 and 8), Non-Tumor Brain (3), SB NPA #1 (4 and 9), Int. NPD, (5 and 11), and SB NPA #2 (10) were aligned using Procedure 1, and the brain tumors: GSC #1 (6 and 13), GSC #2 (14), GSC #3 (15) were aligned using Procedure 2. See fig. S1 for the identity, full H&E images and ROI of the mouse and human samples.

#### Computational Analysis of Nematic Order and Topological Defects

##### Computation of the In-Plane Director Field

The computation of the in-plane director field is based on the procedure in Ref. (*37*). Below, we explain the main steps for completeness.

Let us consider a 2D intensity map *I* (*x, y*), such as a fluorescence image, where (*x, y*) corresponds to the cartesian coordinates in two dimensions. The resolution in the two cartesian directions are considered to be equal. In this work, the resolution was set to 1 *μm*. Next, a Gaussian filter with standard deviation *σ*_1_ was used on *I* (*x, y*). This step eliminated small-wavelength fluctuations in the intensity map. Finally, for every position (*x, y*), we computed the gradients of the intensity map (*∂*_*x*_ *I, ∂*_*y*_ *I*) and constructed a structure matrix *M*_2_, which takes the form

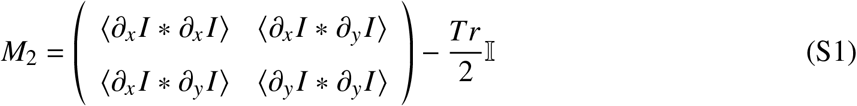

where 𝕀 is the identity matrix. The variable *Tr* = *∂*_*x*_ *I* * *∂*_*x*_ *I* + *∂*_*y*_ *I* * *∂*_*y*_ *I*, and therefore the matrix *M*_2_ is traceless. The symbol ⟨·⟩ denotes a spatial average over a surrounding region at the position (*x, y*). In practice, we used a second Gaussian filter with standard deviation *σ*_2_.

For every position (*x, y*), the matrix *M*_2_ is diagonalized. Then, the eigenvector with the smallest eigenvalue was defined as the in-plane director field **n**_2_(*x, y*). By definition, the in-plane director field **n**_2_(*x, y*) is a unit vector (i.e. |**n**_2_(*x, y*) | = 1). Besides, by definition **n**_2_ and its opposite are equivalent, meaning that **n**_2_ → −**n**_2_ are indistinguishable.

In conclusion, this method computes the in-plane director field **n**_2_ from an intensity map *I* (*x, y*) and two input parameters *σ*_1_, and *σ*_2_. In this work, the value of *σ*_1_ = 1 *μ*m was the same as the image resolution, and the value of *σ*_2_ = 6 *μ*m.

#### Computation of the 3D Director Field

The computation of the 3D director field is based on a generalization of the procedure in Ref. (*37*) from two to three dimensions. Below, we explain the main steps to determine the 3D director field from a 3D intensity map.

Let us consider a 3D intensity map *I* (*x, y, z*), such as a z-stack of fluorescence images, where (*x, y, z*) corresponds to the Cartesian coordinates in three dimensions. Besides, the resolution in the three Cartesian directions is considered to be the same. In this work, the resolution was set to 1 *μm*. Next, a Gaussian filter with standard deviation *σ*_1_ was used on *I* (*x, y, z*). This step eliminated small-wavelength fluctuations in the intensity map. Finally, for every position (*x, y, z*), we computed the gradients of the intensity map (*∂*_*x*_ *I, ∂*_*y*_ *I, ∂*_*z*_ *I*) and constructed a structure matrix *M*_3_, which takes the form

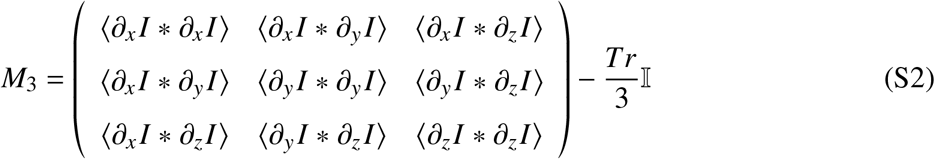

where 𝕀 is the identity matrix. The variable *Tr* = *∂*_*x*_ *I* * *∂*_*x*_ *I* + *∂*_*y*_ *I* * *∂*_*y*_ *I* + *∂*_*z*_ *I* * *∂*_*z*_ *I*, and therefore the matrix *M*_3_ is traceless. The symbol ⟨·⟩ denotes a spatial average over a surrounding region at the position (*x, y, z*). In practice, we used a second Gaussian filter with standard deviation *σ*_2_.

For every position (*x, y, z*), the matrix *M*_3_ was diagonalized. Then, the eigenvector with the smallest eigenvalue was defined as the director field **n**_3_(*x, y, z*). By definition the director field **n**_3_(*x, y, z*) was a unit vector (i.e. |**n**_3_(*x, y, z*) | = 1). Besides, by definition **n**_3_ and its opposite are equivalent, meaning that **n**_3_ → −**n**_3_ are indistinguishable. Note that the **n**_3_ includes the orientation in the (out-of-plane) *z* direction, unlike to the in-plane director field **n**_2_ that was described in the previous section.

In conclusion, the method computes a 3D director field **n**_3_(*x, y, z*) from an 3D intensity map *I* (*x, y, z*), and two input parameter *σ*_1_, and *σ*_2_. In this work, the value of *σ*_1_ = 1 *μ*m was the same as the image resolution, and the value of *σ*_2_ = 6 *μ*m.

##### Computation of 2D Nematic Order Parameters

Below, we explain the procedure to compute from a 3D intensity field *I* (*x, y, z*), two variables that are linked to the nematic order parameter and in the main text, we named 2D nematic order parameter.

For a fixed *z* = *z*_0_ plane, we computed the in-plane director field **n**_2_ from the intensity map *I* (*x, y, z* = *z*_0_), using the method that was described in “Computation of the In-Plane Director Field.” We used this approach because the resolution of H&E sections is lower in the *z* direction than in the *x* and *y* directions. The resolution in *x* and *y* was set to 1 *μ*m. The resolution in *z* was approximately set to a fixed value of 6 *μ*m because the spacing between two consecutive H&E sections was estimated to be 5 *μ*m but roughly 5 of every 6 sections were analyzed, meaning that the average spacing between analyzed section was 6 *μ*m. From the in-plane director field **n**_2_(*x, y, z* = *z*_0_), we determined the phase *ϕ*(*x, y, z* = *z*_0_) as **n**_2_(*x, y, z* = *z*_0_) = (cos(*ϕ*(*x, y, z* = *z*_0_)), sin(*ϕ*(*x, y, z* = *z*_0_))), where the phase *ϕ*(*x, y, z* = *z*_0_) is the angle between the in-plane director field **n**_2_(*x, y, z* = *z*_0_) and the x-axis. Then, the phase *ϕ*(*x, y, z* = *z*_0_) was binned by a factor 6 in the *x* and *y* directions to match the resolution in *x, y, z*. Therefore after this transformation, a unit pixel of *ϕ* corresponds to 6 *μ*m. Finally, repeating these operations for every *z* plane, leads to a 3D map of the phase *ϕ*(*x, y, z*).

Next, we computed the 2D nematic order parameter *S*_2*d*_ (*ℓ*) as

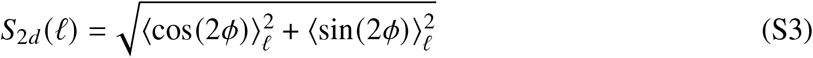

where the symbol ⟨·⟩ denotes a spatial average in the surrounding region of the position (*x, y, z*). In the main text, we computed (S3), in two types of averaging regions: a square in the *x* − *y* plane and a box. In practice, for the former region, we used a 2D square filter, which was computed using the function imboxfilt from Matlab with a filterSize of *ℓ*. Besides, for the latter region, we used a 3D box filter, which was computed using the function imboxfilt3 from Matlab with a filterSize of *ℓ*. Note that the two regions are characterized by a single length scale *ℓ* that is named coarse-graining length. Besides, note that the 2D nematic order parameter that is computed in squares compares the alignment of the in-plane director field within a single image plane, whereas the 2D nematic order parameter that is computed in boxes includes the in-plane director field from other image planes. Finally, *S*_2*d*_ (*ℓ*) was averaged for all positions in the region of interest.

In the case of imaging data from light sheet microscopy, the resolution in *x, y* and *z* was equal and set to 1 *μ*m. To compute the 2D nematic order parameters *S*_2*d*_ (*ℓ*), the phase *ϕ* was binned by a factor 6 in the three cartesian directions. After this transformation a unit pixel corresponds to 6 *μ*m. This allowed to compared the curves of the 2D nematic order parameters with those obtained from 3D reconstructions from sequential H&E sections.

Finally, the range of coarse-graining length studied in experiments was set as follows: the minimal coarse-graining length is 6 *μ*m, which corresponds to one pixel after binning. The maximal coarse-graining length was set by the smallest dimension of a region of interest.

This procedure was used to compute the 2D nematic order parameter shown in Fig. 2, C-F, and figs. S3-S6.

##### Spatial Dependence of the Nematic Order Parameter

A nematic order parameter, such as *S*_2*d*_ (*ℓ*) in “Computation of 2D Nematic Order Parameters” can present three types of scalings with the coarse-graining length *ℓ*. Long-ranged nematic order is when the nematic order parameter is independent of *ℓ, S*(*ℓ*) ∼ *constant*. Quasi-long ranged nematic order is when the nematic order parameter decays as a power law with *ℓ, S*(*ℓ*) *∼ ℓ*^−*α*^, where *α* is the decay exponent. Finally, short-ranged nematic order is when the nematic order parameter decays exponentially with *ℓ, S*(*ℓ*) ∼ exp (−*ℓ*/*λ*), where *λ* is the decay length. Although in statistical physics, these definitions apply in the limit where the coarse-graining length approaches infinity *ℓ* → ∞, here we roughly use the same definitions to describe our experimental system, which has a large cut-off scale set by the system size.

It is instructive to revisit the scaling of the nematic order parameter *S*_2*d*_ (*ℓ*) in Eq. (S3) for a random distribution of orientations. For more details, we refer to (*7*). Let us consider that the phase *ϕ* is a set of independent random variables that are uniformly distributed between 0 and *π*. Then, the nematic order parameter scales as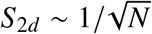, where *N* is the number of independent random variables in the surrounding region. Since in images, the density of pixels is constant, the number *N* is related to the coarse-graining length as *N* ∼ *l*^*d*^, where *d* is the dimension of the surrounding region (*d* = 2 for squares and *d* = 3 for boxes). Therefore, the nematic order parameter scales as *S*_2*d*_ (*ℓ*) ∼ *ℓ*^−*d*/2^ for a random case. The limit of the random case is shown in Fig. 2, and figs. S3-S6 as a black dashed curve.

##### Fitting Procedure of the Curves of 2D Nematic Order Parameter

The procedure to fit the curves of 2D nematic order parameter that were obtained by the procedure explained in “Computation of the 2D Nematic Order Parameters” is the following: For each curve of the 2D nematic order parameter, *S*_2*d*_ (*ℓ*), the power law with a deviation term *f* (*ℓ*) = (*a*−*c*)(*ℓ*/*ℓ*_*c*_)^*b*^+*c* was fitted, where *ℓ* corresponds to the coarse graining length, which in our case corresponds to the length of either a square or a box. The parameter *ℓ*_*c*_ = 6 *μ*m is set to the smallest coarse-graining length. The fitting parameter *a* was constrained in the range of 0.995 to 1.005 because by definition of the nematic order parameter *S*_2*d*_ (*ℓ* = 1*px*) = 1. The other fitting parameter *b* and *c* were unconstrained. The exponent shown in Fig. 2 corresponds to *b* and the asymptotic nematic order at large length scales shown in fig. S7 corresponds to *c*.

##### Computation of the 3D Nematic Order Parameter

From the 3D director field **n**_3_ obtained by the method described in “Computation of the 3D Director Field,” we computed the 3D nematic order parameter as follows:

1. For every position (*x, y, z*), compute a nematic tensor *q* from the component of the director field **n**_3_ = (*n*_*x*_, *n*_*y*_, *n*_*z*_). The nematic tensor *q* takes the form where 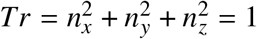 because the director field **n**_3_ is a unit vector (i.e. |**n**_3_| = 1).
2. For every position (*x, y, z*), compute the coarse-grained nematic tensor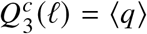, where the symbol ⟨·⟩ denotes a spatial average in the surrounding region of the position (*x, y, z*). In practice, we used a box filter, which was computed using the function imboxfilt3 from Matlab with a filterSize of *ℓ*. The parameter *ℓ* is the coarse-graining length.
3. For every position (*x, y, z*), diagonalize the coarse-grained nematic tensor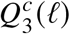. Since by definition, the nematic tensor is uniaxial, the diagonalised form of 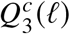 reads

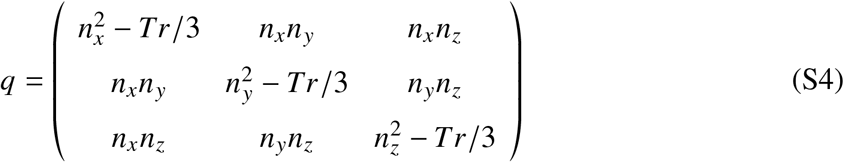

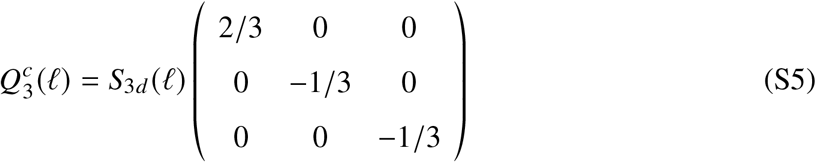

where *S*_3*d*_ (*ℓ*) is the 3D nematic order parameter.

This procedure was applied only to the case of imaging data from light sheet microscopy. Finally, we compared the curves of 2D nematic order parameters that were computed in boxes to the 3D nematic order parameter *S*_3*d*_ (*ℓ*), Fig. 1, I and J. As it is expected the two nematic order parameters decay with the coarse-graining length *ℓ*, Fig. 1, I and J. Next, we fitted a power law with exponent *b* and a deviation term *c* as explained in “Fitting Procedure of the Curves of 2D Nematic Order Parameter.” The average decay exponent of *S*_3*d*_ is −0.29 ± 0.06 (mean±SD, *n* = 10), was similar to the average decay exponent of *S*_2*d*_ is −0.36 ± 0.10. Besides the average values of *c* was also similar between the two cases: 0.09 ± 0.14 for *S*_3*d*_ and 0.08 ± 0.17 for *S*_2*d*_. Therefore, this suggests the scaling of the 2D nematic order parameter was a good metric of the 3D nematic alignment.

##### Computation of Low Nematic Order Isosurfaces

The coarse-grained nematic tensor 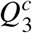 was computed as explained in the above Section, except that the director field **n**_3_ was not binned by a factor 6. The coarse graining length was set to *ℓ* = 31 *μ*m.

To visualize the coarse-grained nematic tensor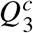, we used the software Open-ViewMin at https://gitlab.com/open-viewmin/open-viewmin.gitlab.io. The green ellipsoids in Fig. 3 and figs. S8-S10 represent the director field associated with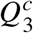, which is uniaxial by definition. The purple surfaces in the same figures correspond to low nematic order isosurfaces. These were also computed using the software Open-ViewMin. In short, the surfaces are defined by the condition that the largest eigenvalue of 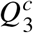 is equal to a number *λ*. In our case, this number was between *λ* = 0.2 and 0.23. The white outlines in the same figures correspond to cuts of a low nematic order surface with a *z* plane. We checked that the shape of the isosurface and the director field were qualitatively similar upon small changes in both the parameter *λ* and *ℓ*.

Finally, in the same figures, the pink regions roughly represent vessels. These regions were determined by binarising images based on an intensity threshold that was adjusted for each region of interest independently.

##### Analysis of +1/2 Topological Defect Motion

Here, we analyzed time-lapse images of cultures for up to 16 h to 77 h post-confluency. For each time point, the two-dimensional director field **n**_2*d*_ was computed using the method explained in “Computation of the In-Plane Director Field” using a value of *σ*_2_ = 21 *μ*m. Half-integer topological defects were identified as regions where the 2*d* nematic order parameter *S*_2*d*_ < 0.2 and the winding number of the director field is ±1/2. Here *S*_2*d*_ was computed using a Gaussian filter with standard deviation of 7 *μ*m. The polarity **p** of +1/2 topological defects is computed using Eq. (5) in Ref. (*38*). If two topological defect trajectories are closer than 150 *μ*m, both defects are eliminated to exclude motion due to defect interactions.

Tracking of +1/2 topological defects was performed using the Python module trackpy with the parameter ‘search range’ = 83 *μ*m and the parameter ‘memory’= 50 frames, (*39*). The second parameter is between 8 h and 38 h depending on the movie. If a defect trajectory consists of less than 10 frames or its length is shorter than 30 *μ*m, it is eliminated.

Finally, we analyzed the motion of the remaining +1/2 topological defects, *N* = 172, see fig. S17A. For each of trajectory, we computed the angle difference Δ*θ* between the mean defect polarization, and the displacement vector from the last time point to the first time point, see fig. S17B. The mean defect polarization was computed as the mean of **p** over a trajectory. The motion of the defect was classified as head-to-tail, if −45° < Δ*θ* < 45°, tail-to-head, if 225° < Δ*θ* < 135°, and ‘other’ in any other case, see fig. S17C. The results in fig. S17D show that +1/2 defect motion was predominantly tail-to-head.

**Caption for Movie S1: (LSM Z-Stack)** Raw light-sheet microscopy *z*-stack image of GFP-tagged (green) glioma NPD in vivo tumor. *Z*-planes are 1 *μ*m apart. The cell orientations form a trefoil-like shape in 2D which propagates in the *z*-direction. The -1/2 defect is analyzed in detail in Fig. 3 and fig. S10.

**Caption for Movie S2: (Omega Vector Rotations)** Rotating view of nematic defects in the cleared intracranial NPD mouse glioma region shown in Fig. 3, with analysis of defect winding geometry. Transparent purple surfaces are isosurfaces of nematic order bounding low-order regions, including topological defects, as in Fig. 3. Pink surfaces indicate blood vessels. At a sampled subset of points on the low-order surfaces, the nematic director field is plotted as teal cylinders at nearby points located on a small circuit, which encircles the associated point on the defect provided the surface is tube-like there. Double-headed arrows (shaded green when twist-type to yellow when wedge-type) show the rotation vector, Ω (with sign ambiguity) at those defect points, computed from the normal to the plane of rotation of the director on the circuit.

**Caption for Movie S3: (X-Y View of Fig 3)** Animation of changing *z*-slice coordinate in a three-dimensional view of the nematic order and defects in the cleared intracranial NPD mouse glioma region shown in Fig. 3. On each slice of constant *z*, gray-scale LSM intensity is plotted along with the inferred nematic director field (teal cylinders). Purple surfaces are isosurfaces of nematic order bounding low-order regions, including topological defects, as in Fig. 3. Pink surfaces indicate blood vessels.

**Caption for Movie S4: (X-Z View of Fig 3)** Animation of changing y-slice coordinate in a three-dimensional view of the nematic order and defects in the cleared intracranial NPD mouse glioma region shown in Fig. 3. On each slice of constant *y*, gray-scale LSM intensity is plotted along with the inferred nematic director field (teal cylinders). Purple surfaces are isosurfaces of nematic order bounding low-order regions, including topological defects, as in Fig. 3. Pink surfaces indicate blood vessels.

**Caption for Movie S5: (05-03-23-Pos3-GFP)** 20x in vitro confocal movie showing formation of trefoils and comets in GFP-tagged (green) glioma NPA adherent cells over 70 hours. Movie analyzed in fig. S16, A-D.

**Caption for Movie S6: (07-12-22-Pos2-GFP)** 20x in vitro confocal movie showing formation of trefoils and comets in GFP-tagged (green) glioma NPA adherent cells over 25 hours.

**Caption for Movie S7: (05-19-23-Pos4-Casp3)** 20x in vitro confocal movie showing formation of trefoils and comets in GFP-tagged (green) glioma NPA adherent cells over 70 hours with apoptosis labeled in red.

**Caption for Movie S8: (05-19-23-Pos9-Casp3)** 20x in vitro confocal movie showing formation of trefoils and comets in GFP-tagged (green) glioma NPA adherent cells over 70 hours with apoptosis labeled in red.

**Figure S1:**
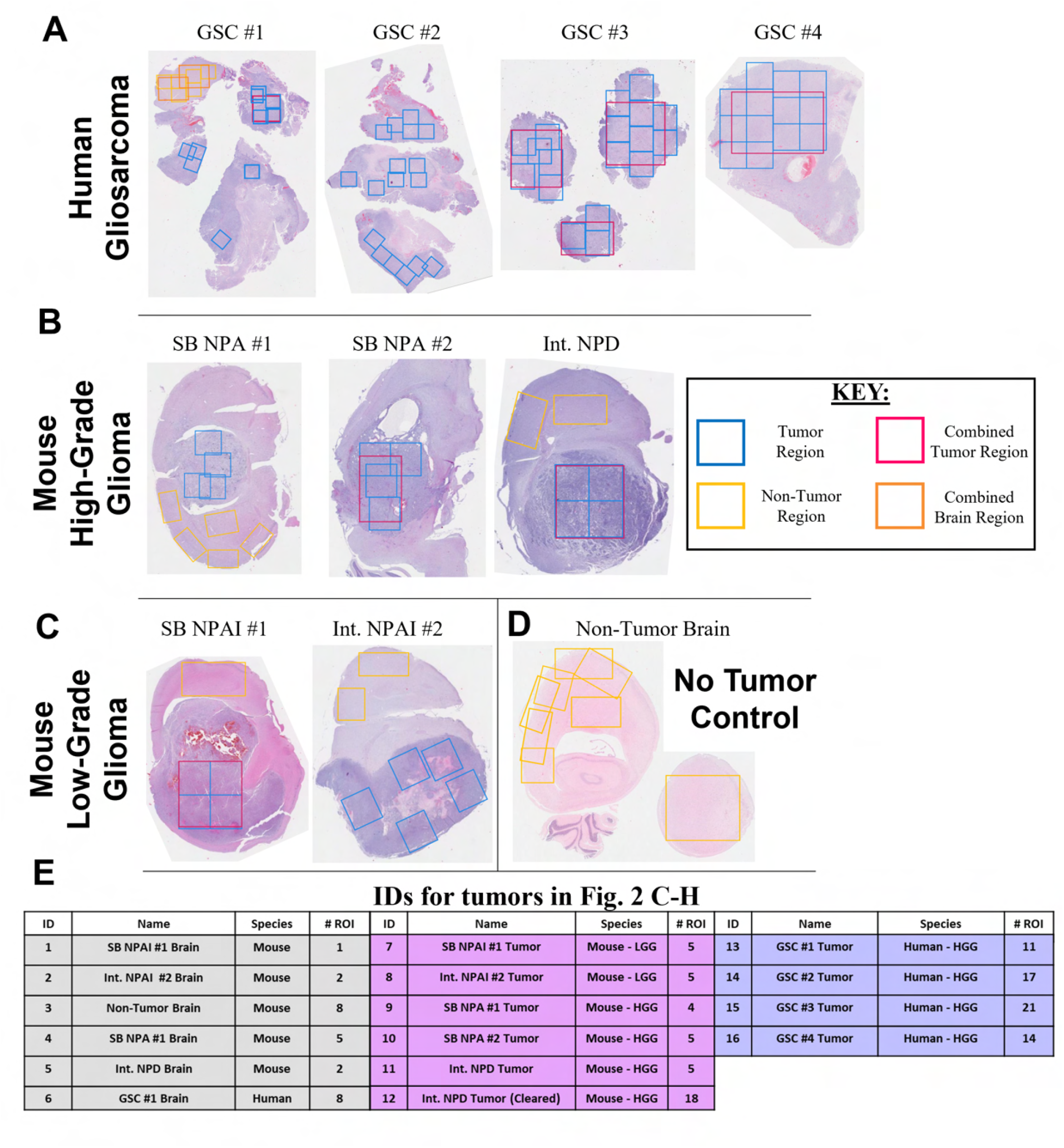
H&E brain samples from human gliosarcoma patients and mouse models. Examples of H&E sections of each brain and tumor analyzed. **(A)** Human gliosarcoma (GSC) patient samples. **(B)** High-grade mouse gliomas type NPA and NPD. **(C)** Low-grade mouse glioma models type NPAI. **(D)** Normal brain control with no tumor. Regions of interests (ROIs) are outlined with colorcoded boxes to illustrate the regions analyzed. Blue are tumor regions, pink are larger combined tumor regions, yellow are non-tumor regions like tumor-adjacent normal brain regions or brain control regions, and orange are larger combined non-tumor regions. **(E)** Corresponding brain/tumor names for ID numbers found in Main Figure 2 with animal species and numbers of ROI listed. Abbreviations for tumor ID names defined in Materials and Methods. The acronym SB stands for Sleeping Beauty and the acronym Int. for intracranial tumor generation methods.

**Figure S2:**
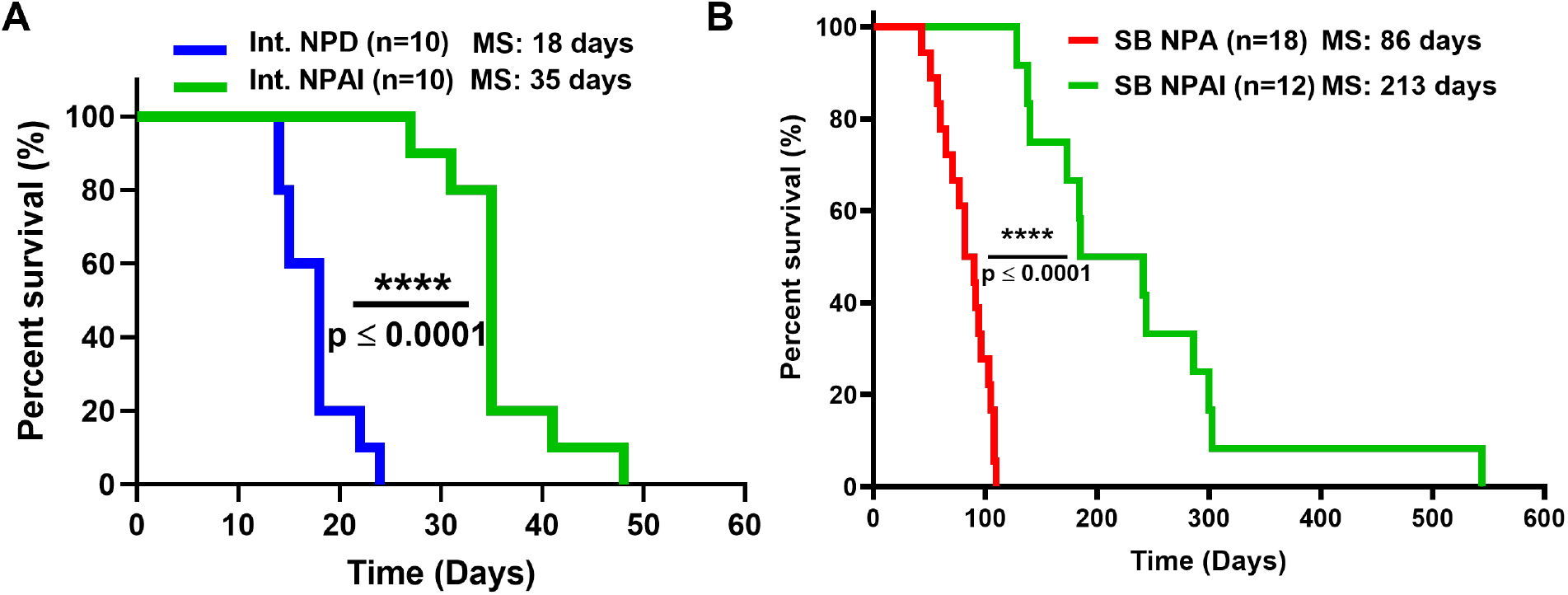
Glioma Survival Curves. **(A)** Kaplan-Meier survival curves for Int. NPD and Int. NPAI mouse tumors (*P* value * * * ≤ 0.0001). **(B)** Kaplan-Meier survival curves for SB NPA and SB NPAI mouse tumors (*P* value * * * ≤ 0.0001). Abbreviations for tumor ID names defined in Materials and Methods. NPA and NPD are high-grade gliomas while NPAI is low-grade. Figure modified from (*18*).

**Figure S3:**
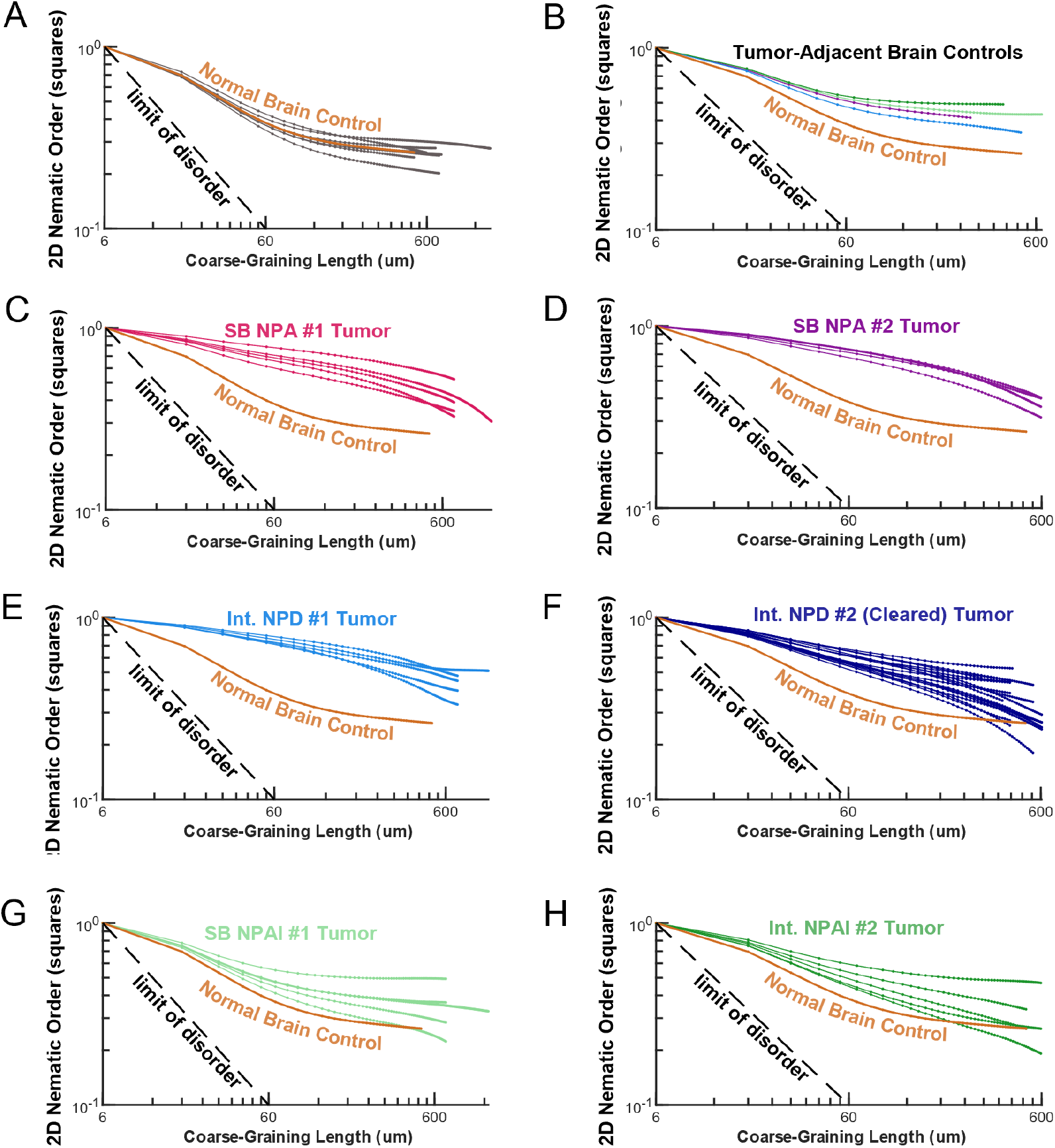
Nematic Order Parameter in Mouse Tumors in 2D. (A-H) 2D Nematic Order Parameter (NOP) as a function of coarse-graining length of the averaging domains (squares) is shown for a variety of mouse tumors of different aggression levels (C-H), paired tumor-adjacent regions (B), and a control normal brain (A). High Grade Gliomas: panels C-F. Low Grade Gliomas: panels G-H. Each tumor or normal brain piece is segmented into several smaller regions which are plotted in the same color. The average NOP of the normal brain control graphed in panel A is shown as the orange solid curve in all panels. The black dashed curve in all panels represents the theoretical limit of a random case *y* = *x*^−1^. Color keys: (A) Control brain containing no tumor - gray. (B) tumor-adjacent brain controls from tumors shown in (D,E,G,H) with matching colors. (C) SB NPA #1 tumor - pink. (D) SB NPA #2 tumor - purple. (E) Int. NPD #1 tumor - light blue. (F) Int. NPD #2 tumor, imaged with clearing - dark blue. (G) SB NPAI #1 tumor - light green. (H) Int. NPAI #2 tumor - dark green. Abbreviations for tumor ID names defined in Materials and Methods. The acronym SB stands for sleeping beauty and the acronym Int. for intracranial.

**Figure S4:**
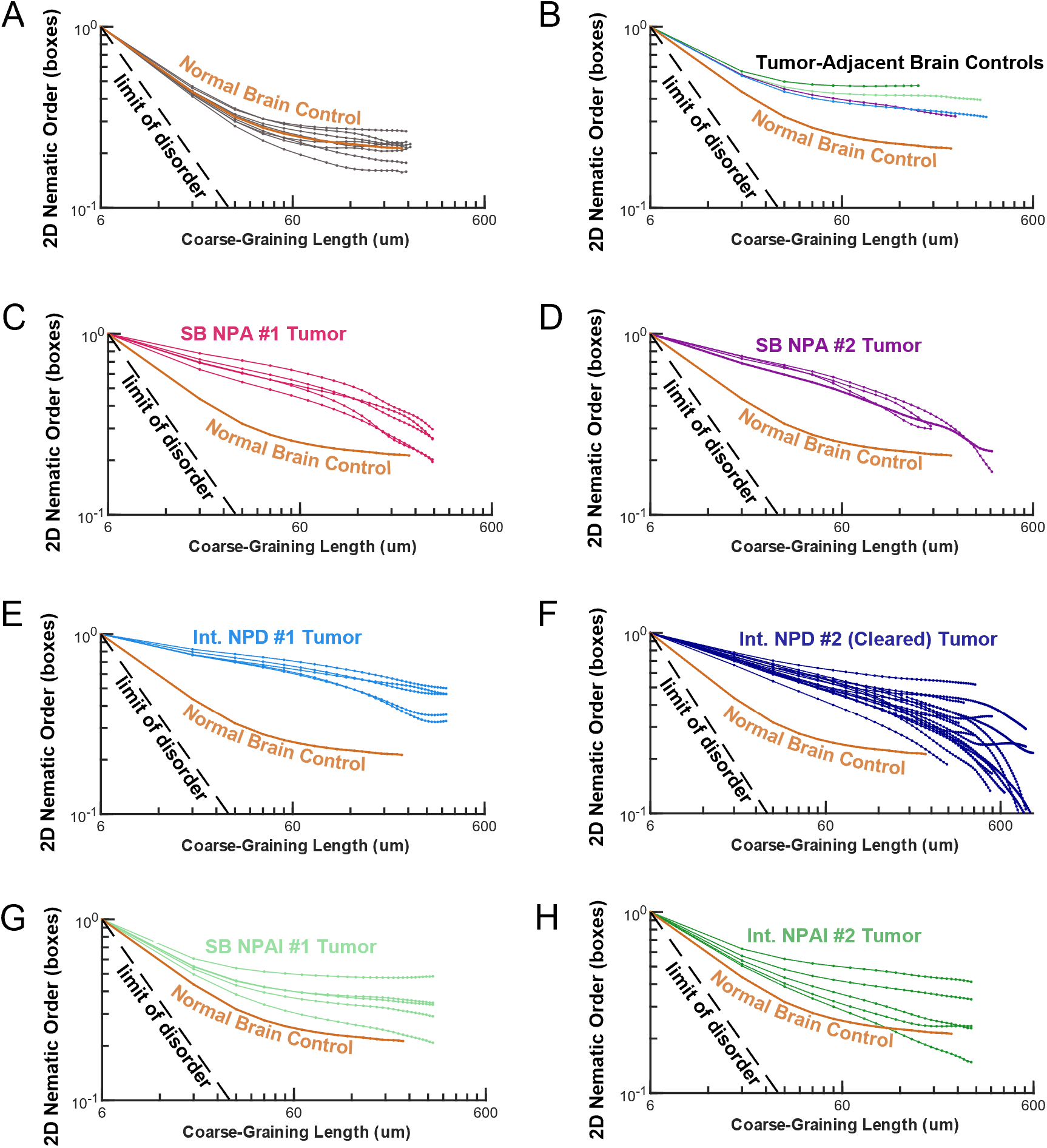
Nematic Order Parameter in Mouse Tumors in 3D. Same as Figure S3, but plotting the 2D Nematic Order Parameter (NOP) as a function of coarse-graining length of the averaging domains in **3D boxes**. Here, the theoretical limit of a random case is *y* = *x*^−1.5^.

**Figure S5:**
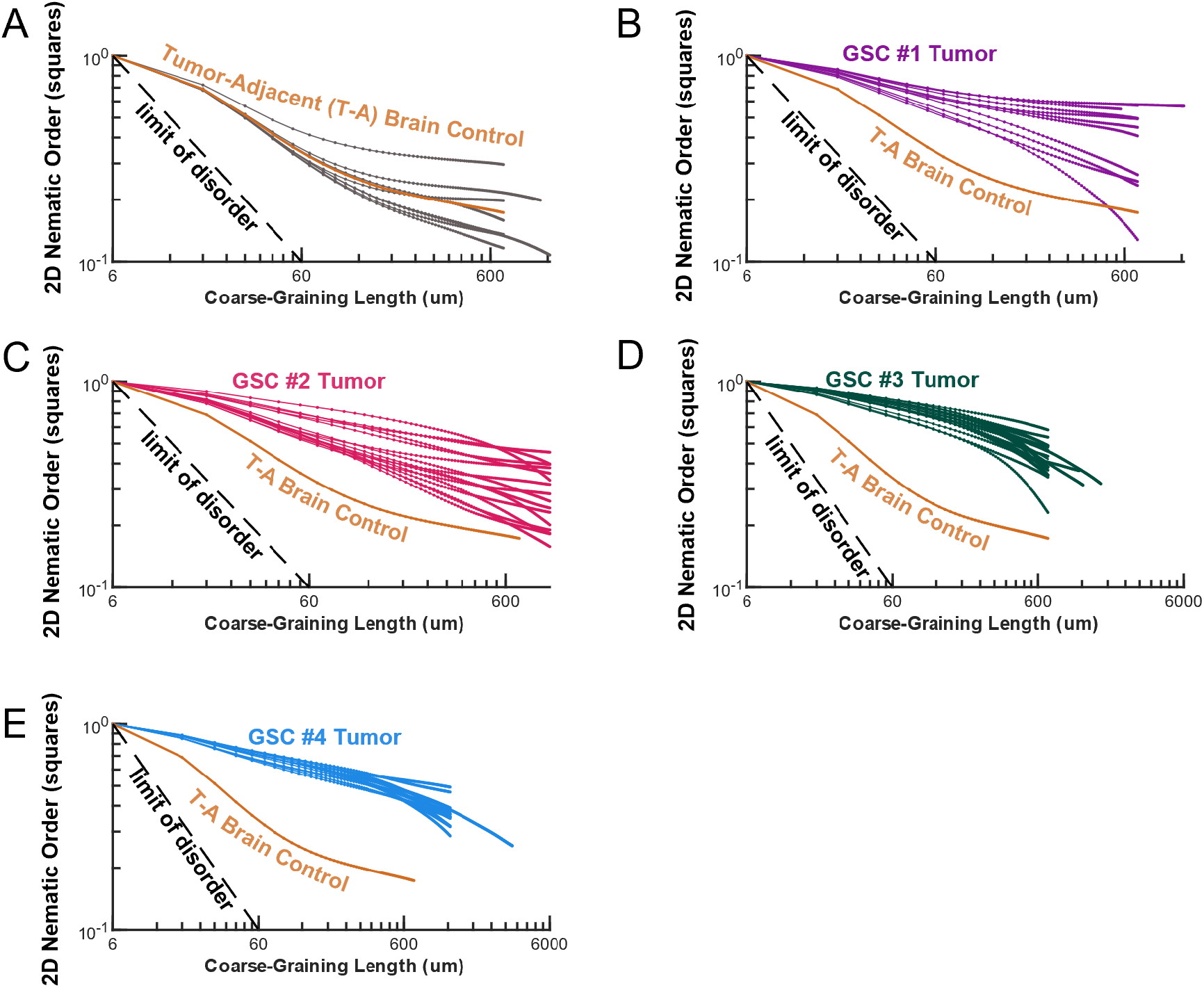
Nematic Order Parameter in Human Gliosarcoma Tumors in 2D. Same as Figure S4, but plotting for a variety of human gliosarcoma (GSC) tumors and a control tumor-adjacent normal brain region. Color keys: (A) GSC #1 tumor-adjacent normal brain control regions - gray. (B) GSC #1 tumor - purple. (C) GSC #2 tumor - pink. (D) GSC #3 tumor - dark green. (D) GSC #2 tumor - blue. Abbreviations for tumor ID names defined in Materials and Methods and S1E.

**Figure S6:**
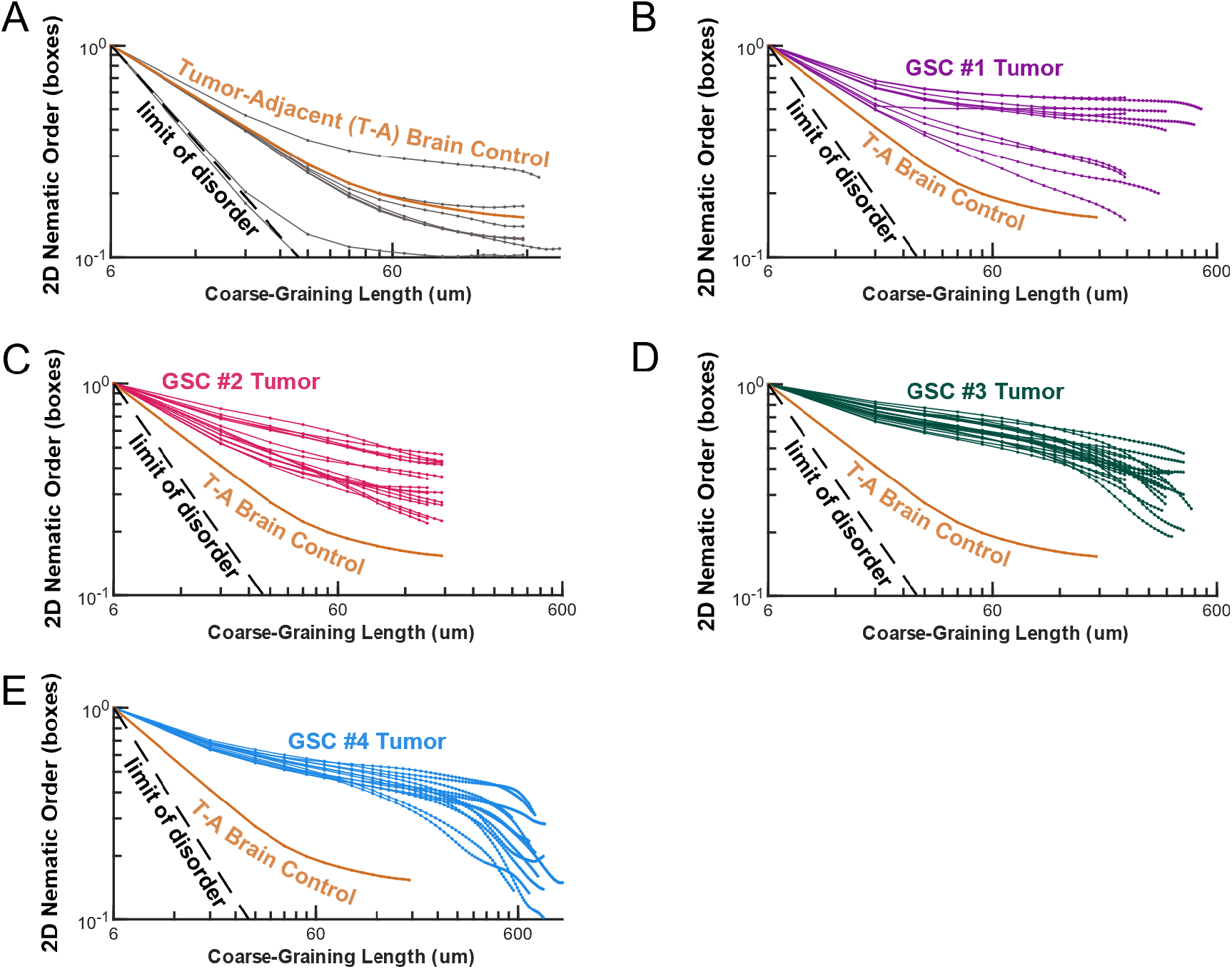
Nematic Order Parameter in Human Gliosarcoma Tumors in 3D. Same as Figures S3 and S5, but plotting the 2D Nematic Order Parameter (NOP) as a function of coarse-graining length of the averaging domains in **3D boxes** in human gliosarcoma samples. Here, the theoretical limit of a random case is *y* = *x*^−1.5^.

**Figure S7:**
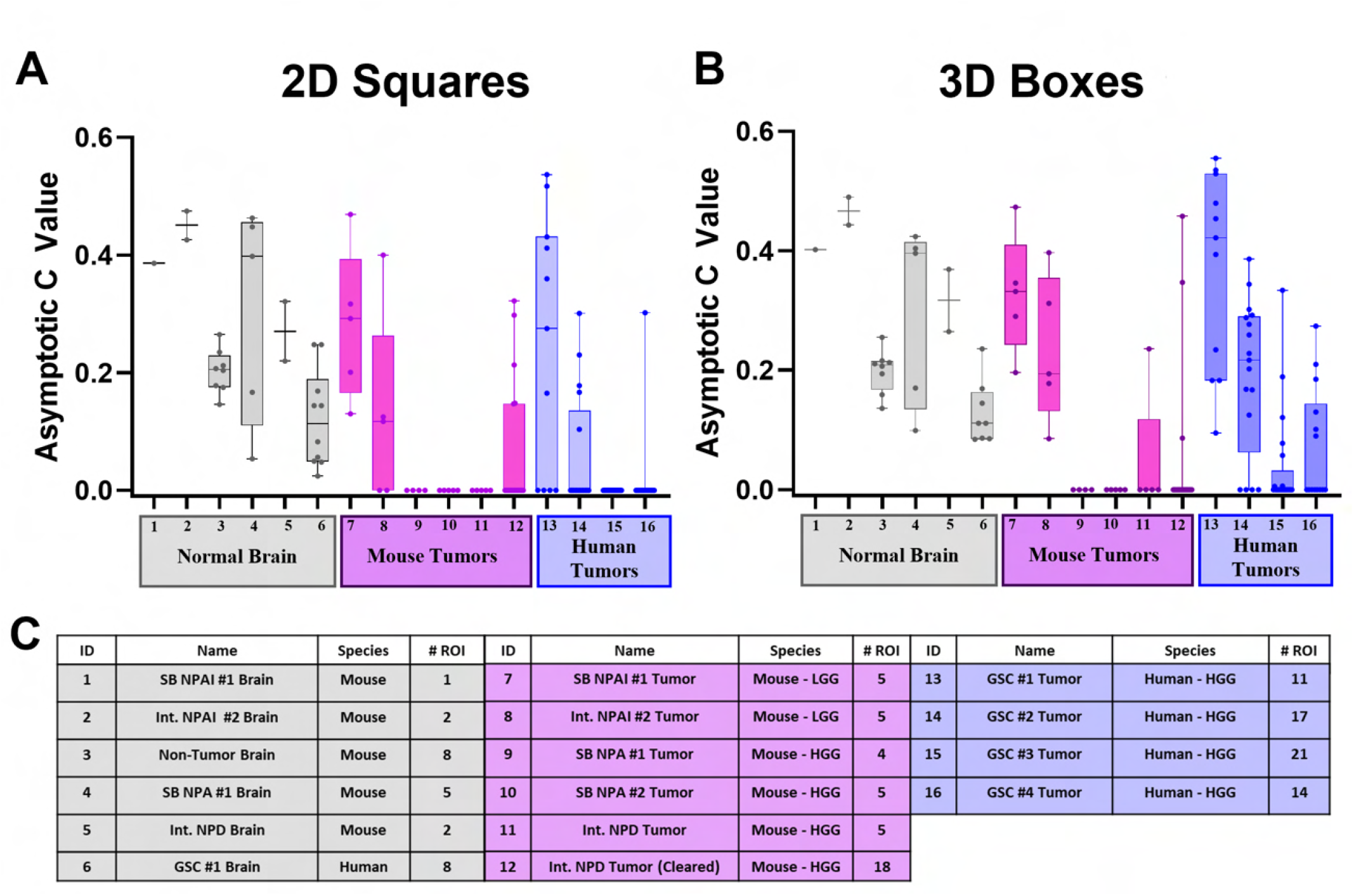
2D and 3D asymptotic level of nematic order at large length scales. (**A-B**) Respectively, 2D and 3D asymptotic *c* values for various tumor and normal brain regions of interest (ROI). For each ROI, the 2D nematic order parameter curves that is computed in either squares or boxes was fitted with a power law: *f* (*x*) = (1 − *c*)*x*^*b*^ + *c* (see Materials and Methods for definitions), respectively. The asymptotic level of nematic order at large length scales corresponds to the fitting parameter *c*. Box and whisker plot shows each ROI’s value of *c* with mean shown as horizontal line. X-axis labels correspond to each tissue listed in the table of panel fig. S1E, shown again in panel (**C**). Shaded areas correspond to the standard deviation over multiple ROIs.

**Figure S8:**
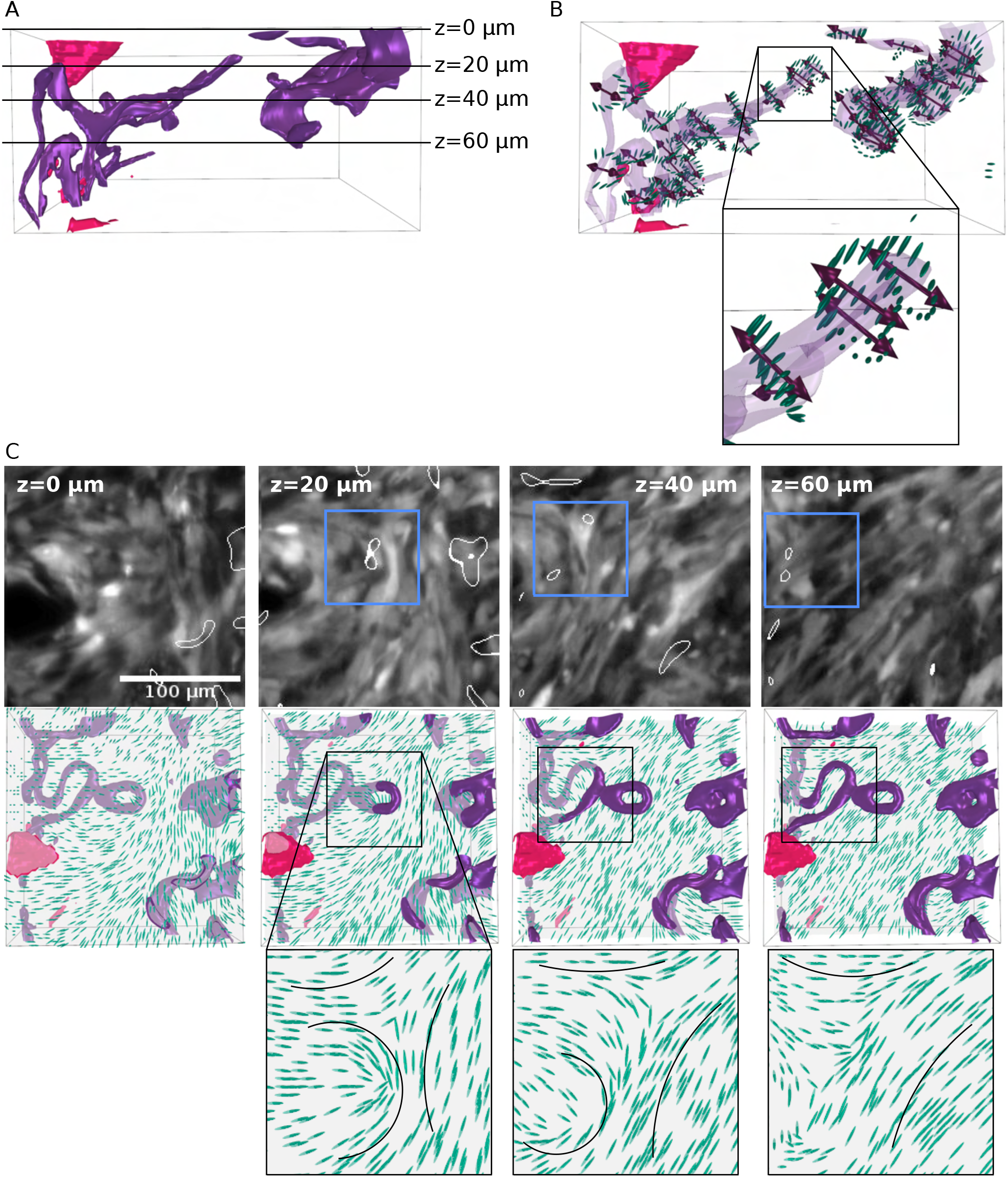
Example of twist-type disclination loop. **(A)** 3D reconstructions of a region from cleared intracranial NPD mouse glioma. Dark purple regions correspond to surfaces with low nematic order. Pink regions outline blood vessels. The black lines label the *z*-planes that are shown in C. (**B**) Detailed side view of the 3D reconstruction in panel A. To facilitate visualization, the dark purple areas in panel A are now translucent purple. The 3D director field in the surrounding purple regions is shown as green ellipsoids and the double-headed red arrows represent the rotation vector, Ω. Below: approximate zoom in of the black square in panel B. (**C**) Top row: Gray-scale LSM images of cleared tumor with blue boxes outlining disclination lines (thin white outline) through the *z*-plane (*z* coordinate indicated in top left corner in white). Middle row: Corresponding 3D director field (green ellipsoids) reconstructions with purple regions corresponding to surfaces with low nematic order and disclination-line locations now outlined with black boxes. Note: the *z*-plane is translucent gray, and thus the features above the *z*-plane appear in a darker color and the features below the *z*-plane in a dimmer color. Bottom row: Zoomed in view of disclination lines traveling through the *z*-plane. In the left-most column, there is no panel as the disclination of interest does not appear at *z* = 0 *μ*m (see panel A). The scale bar is 100 *μ*m.

**Figure S9:**
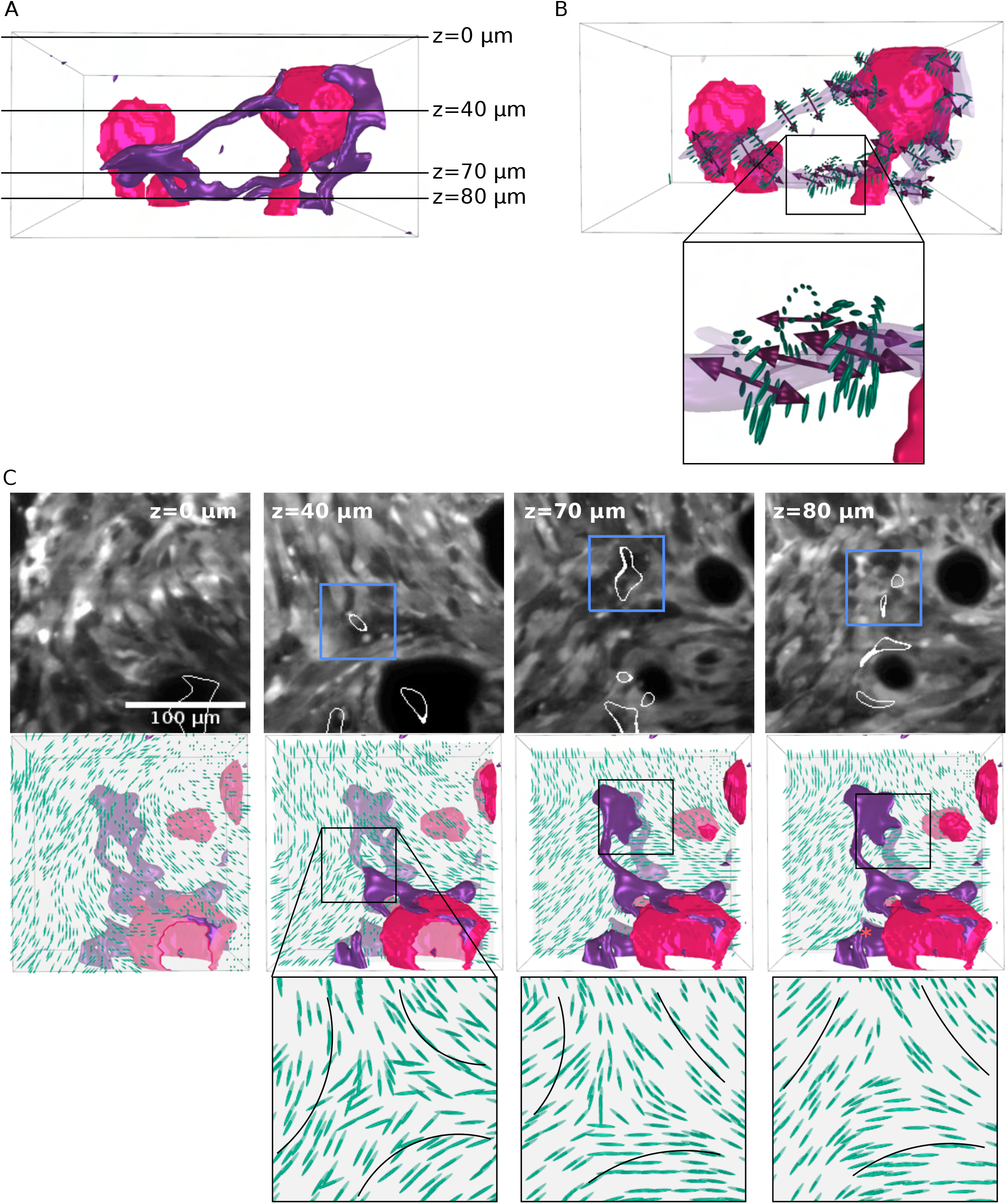
Example of a wedge-type disclination line that starts and ends on a blood vessel. (A) 3D reconstructions of a region from cleared intracranial NPD mouse glioma. Dark purple regions correspond to surfaces with low nematic order. Pink regions outline blood vessels. The black lines label the *z*-planes that are shown in C. (**B**) Detailed side view of the 3D reconstruction in panel A. To facilitate visualization, the dark purple areas in panel A are now translucent purple. The 3D director field in the surrounding purple regions is shown as green ellipsoids and the double-headed red arrows represent the rotation vector, Ω. Below: zoom in of the black square in panel B. (**C**) Top row: Gray-scale LSM images of cleared tumor with blue boxes outlining disclination lines (thin white outline) through the *z*-plane (*z* coordinate indicated in top corner of image). Middle row: Corresponding 3D director field (green ellipsoids) reconstructions with purple regions corresponding to surfaces with low nematic order and disclination-line locations now outlined with black boxes. Note: the *z*-plane is translucent gray, and thus the features above the *z*-plane appear in a darker color and the features below the *z*-plane in a dimmer color. Bottom row: Zoomed in view of disclination lines traveling through the *z*-plane. In the left-most column, there is no panel as the disclination of interest does not appear at *z* = 0 *μ*m (see panel A). The scale bar is 100 *μ*m.

**Figure S10:**
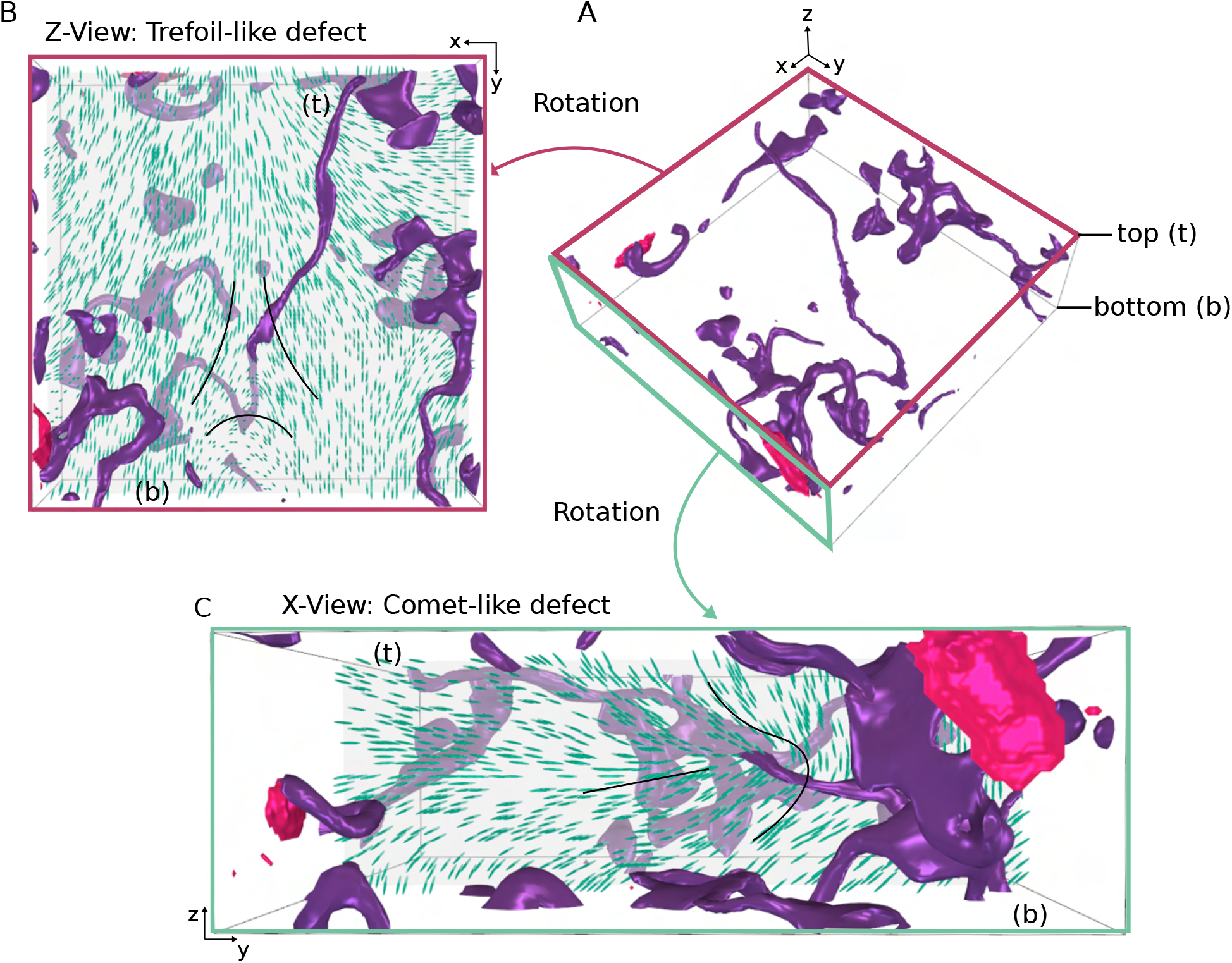
Two cross-sections of a twist-like disclination. (**A**) Same 3D reconstruction as in panel A in Fig. 3. Dark purple regions correspond to surfaces with low nematic order (see Methods). Pink regions roughly outline blood vessels. The top z-plane is labeled (t) and the bottom plane is labeled (b). (**B**) Z-view of the 3D reconstruction in panel A. The red outline corresponds to the red face in panel A. 3D director field is shown as green ellipsoids. In this view, the defect appears as a -1/2 trefoil. (**C**) X-view of the 3D reconstruction in panel A. The green outline corresponds to the green face in panel A. To improve visualization, the scale of panel C was increased by a factor 2 with respect to the other panels. In this view, the defect appears as a +1/2 trefoil. In both panels B and C, the director field around the disclination line is highlighted with black curves. The dimensions of the region in panel A are 300 × 300 × 100 *μ*m^3^.

**Figure S11:**
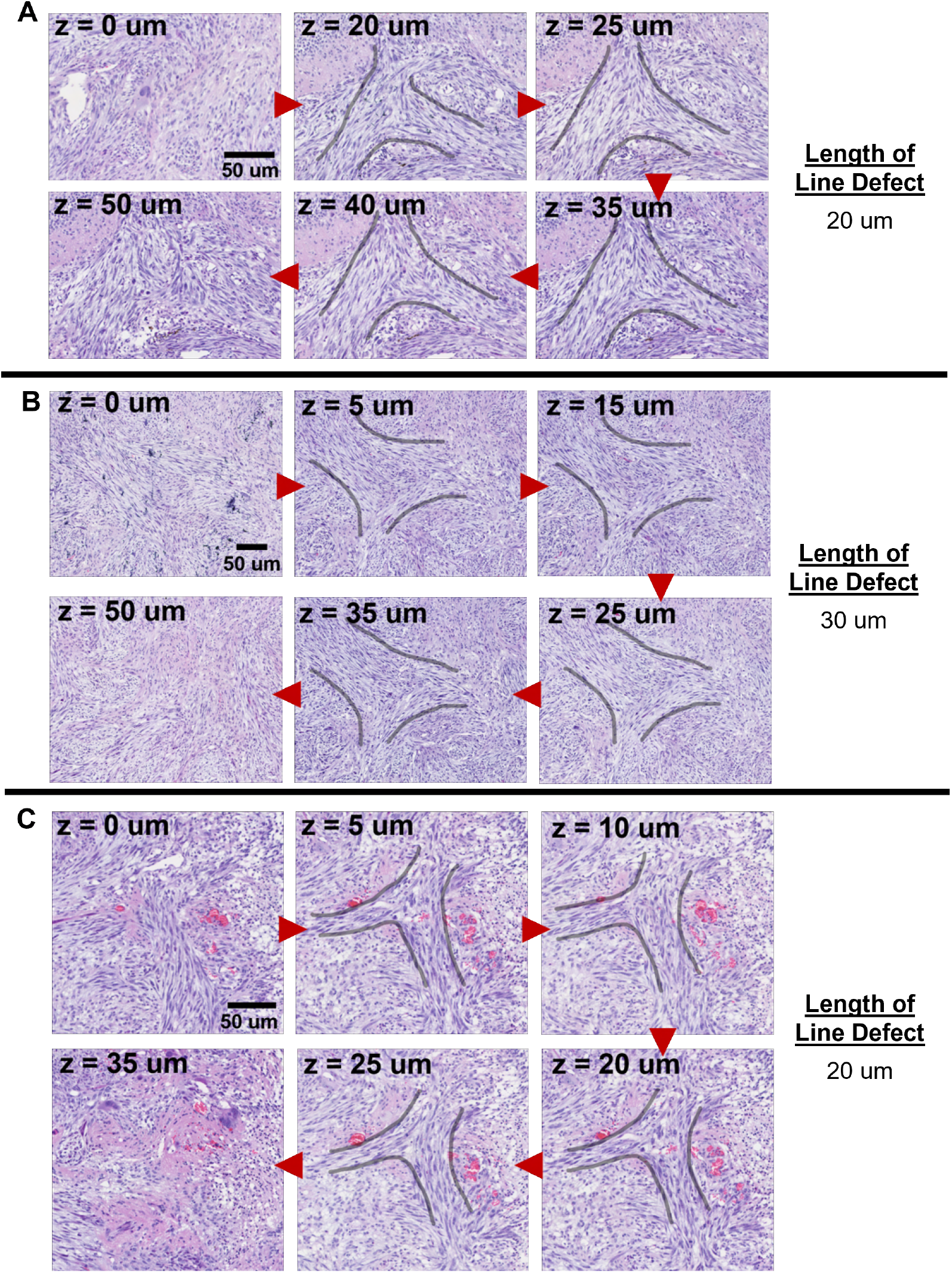
Trefoil Topological Defect Lines in SB NPA Tumor #2. **(A-C)** Three examples of H&E stained serial sections of Sleeping Beauty NPA #2 tumor highlighting trefoil topological defects over several slices. Each image shows the same region within the tumor at different *z* depth, as indicated in the top left corner. Red arrows show progression into the *z*-axis of the tumor. Gray curves outline trefoils. The length of the defect sequences is indicated on the right. Scale bar = 50 *μ*m.

**Figure S12:**
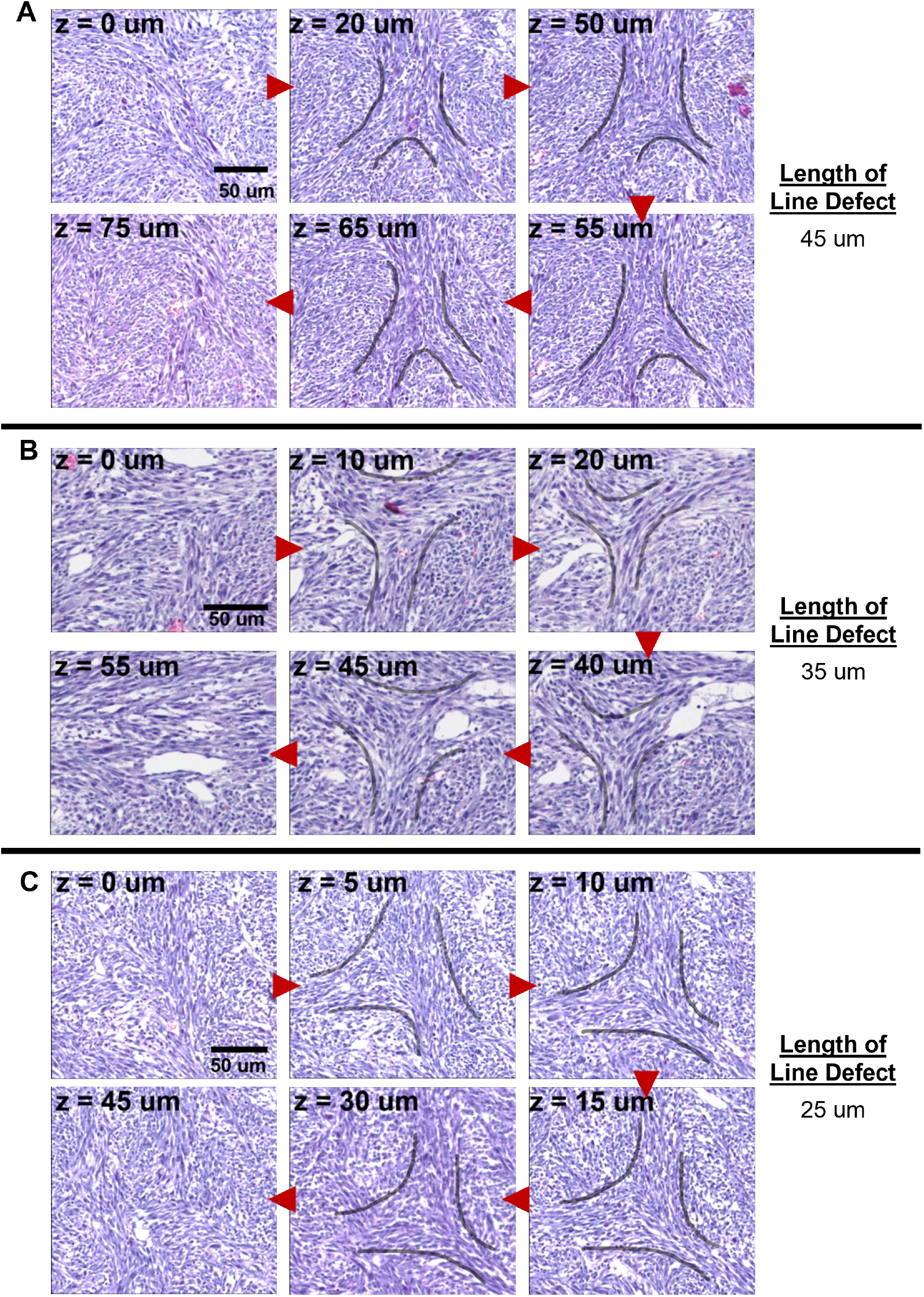
Trefoil Topological Defect Lines in SB NPD Tumor #1. **(A-C)** Three examples of H&E stained serial sections of intracranial NPD #1 tumor highlighting trefoil topological defects over several slices. Each image shows the same region within the tumor at different *z* depth, as indicated in the top left corner. Red arrows show progression into the *z*-axis of the tumor. Gray curves outline trefoils. The length of the defect sequences is indicated on the right. Scale bar = 50 *μ*m.

**Figure S13:**
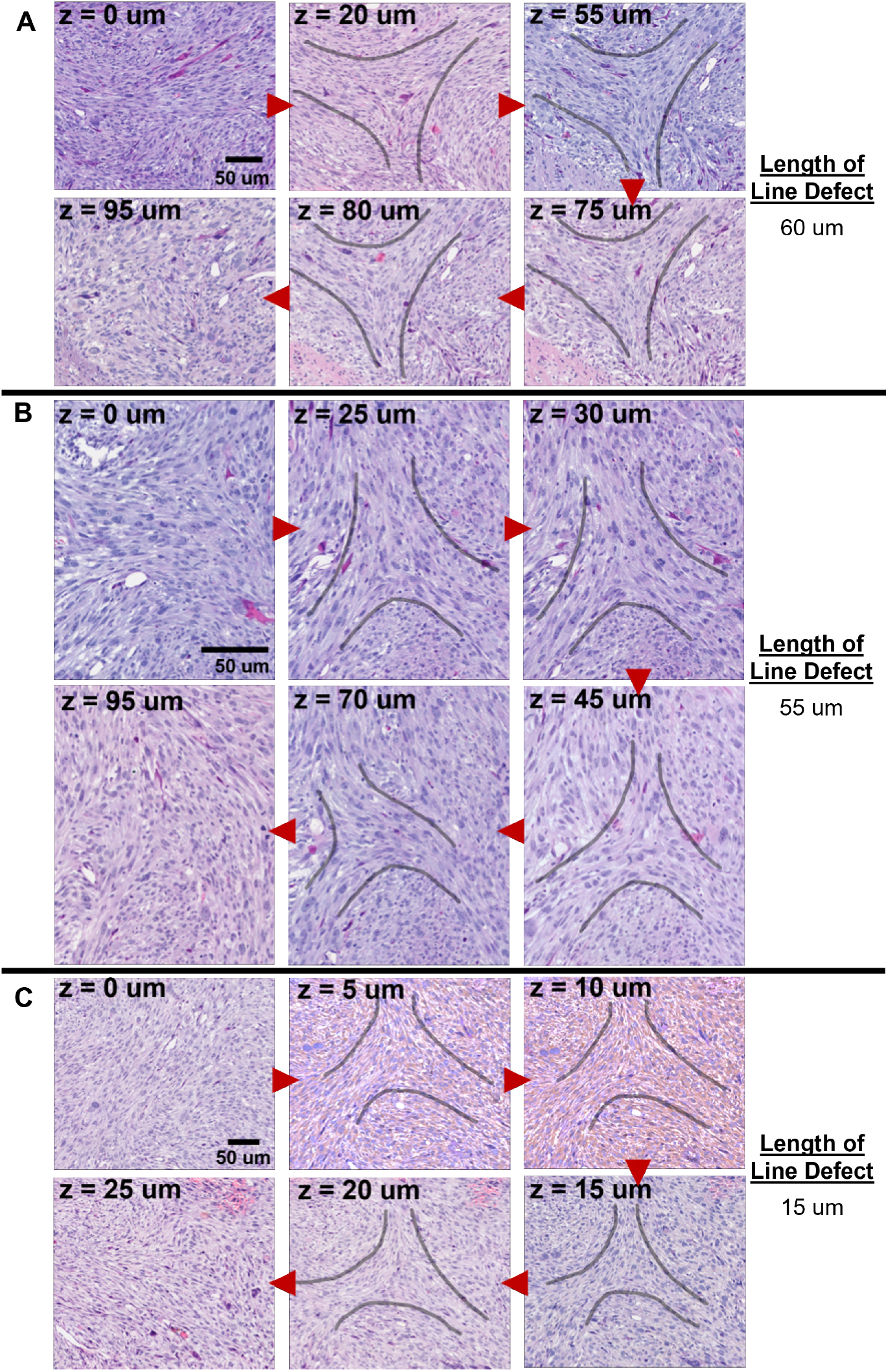
Trefoil Topological Defect Lines in SB NPA Tumor #1. **(A-C)** Three examples of H&E stained serial sections of Sleeping Beauty NPA #1 tumor highlighting trefoil topological defects over several slices. Each image shows the same region within the tumor at different z depth, as indicated in the top left corner. Red arrows show progression into the z-axis of the tumor. Gray curves outline trefoils. The length of the defect sequences is indicated on the right. Scale bar = 50 *μ*m.

**Figure S14:**
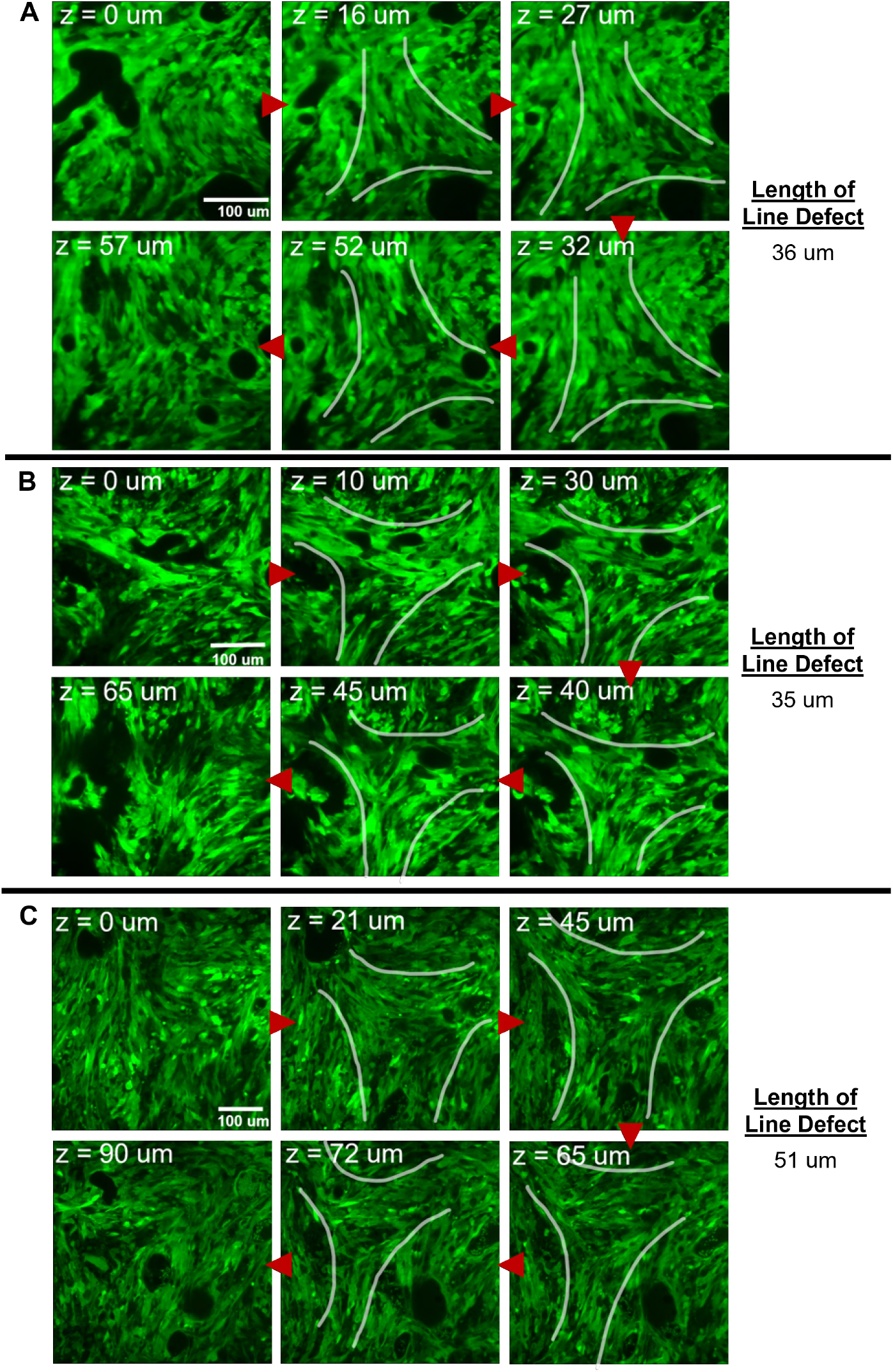
Trefoil Topological Defect Lines in Cleared Intracranial NPD Tumor. **(A-C)** Three examples of LSM imaged cleared intracranial NPD tumors (*n* = 3) highlighting trefoil topological defects over several slices. Each image shows the same region within the tumor at different *z* depth, as indicated in the top left corner. Red arrows show progression into the *z*-axis of the tumor. Gray curves outline trefoils. The length of the defect sequences is indicated on the right. Scale bar = 100 *μ*m.

**Figure S15:**
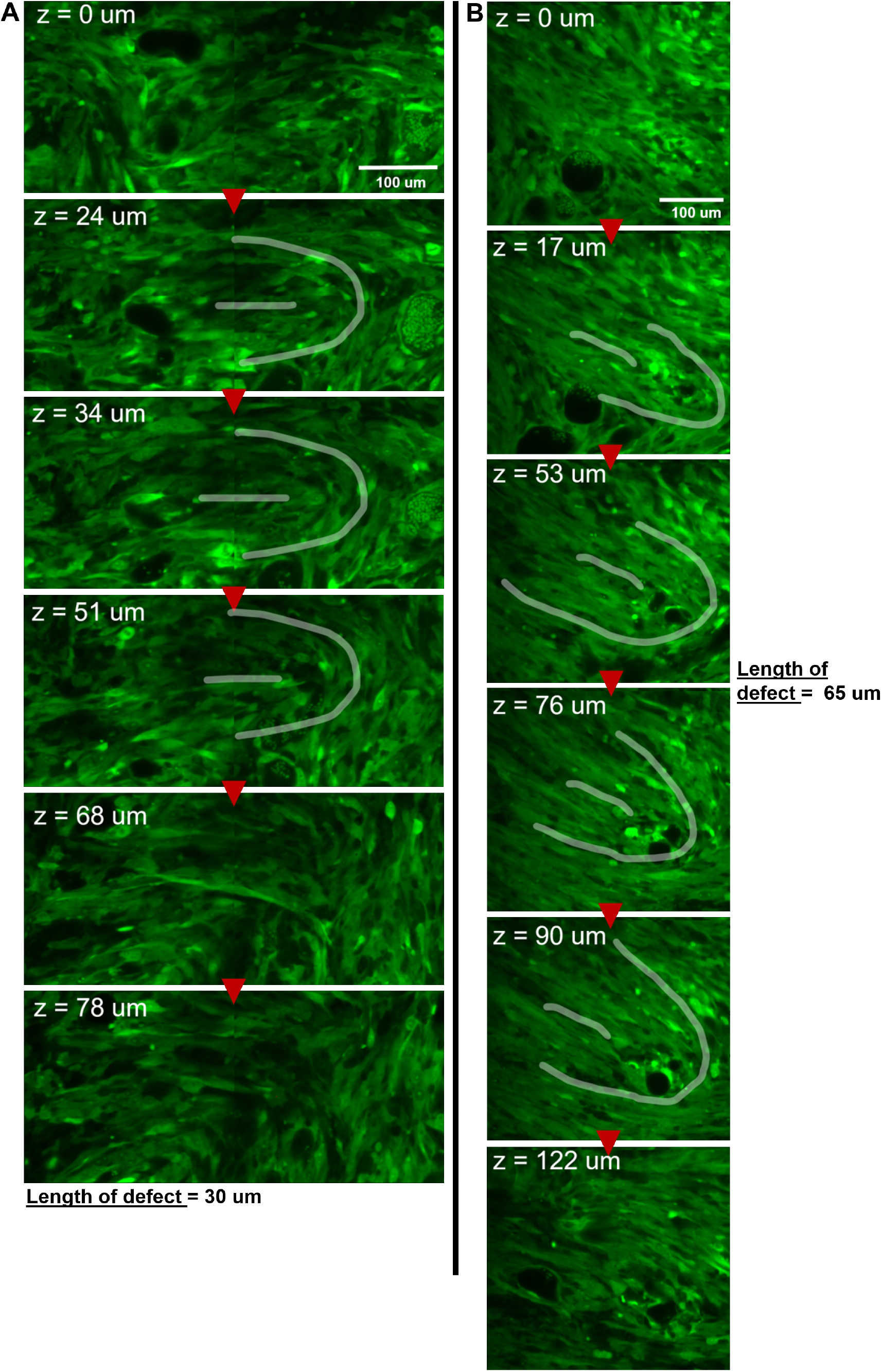
Comet Topological Defect Lines in Cleared Intracranial NPD Tumor. **(A-B)** Two examples of LSM imaged cleared intracranial NPD tumors highlighting comet topological defects over several slices. Each image shows the same region within the tumor at different *z* depth, as indicated in the top left corner. Red arrows show progression into the *z*-axis of the tumor. Gray curves outline comets. The length of the defect sequences is indicated. Scale bar = 100 *μ*m.

**Figure S16:**
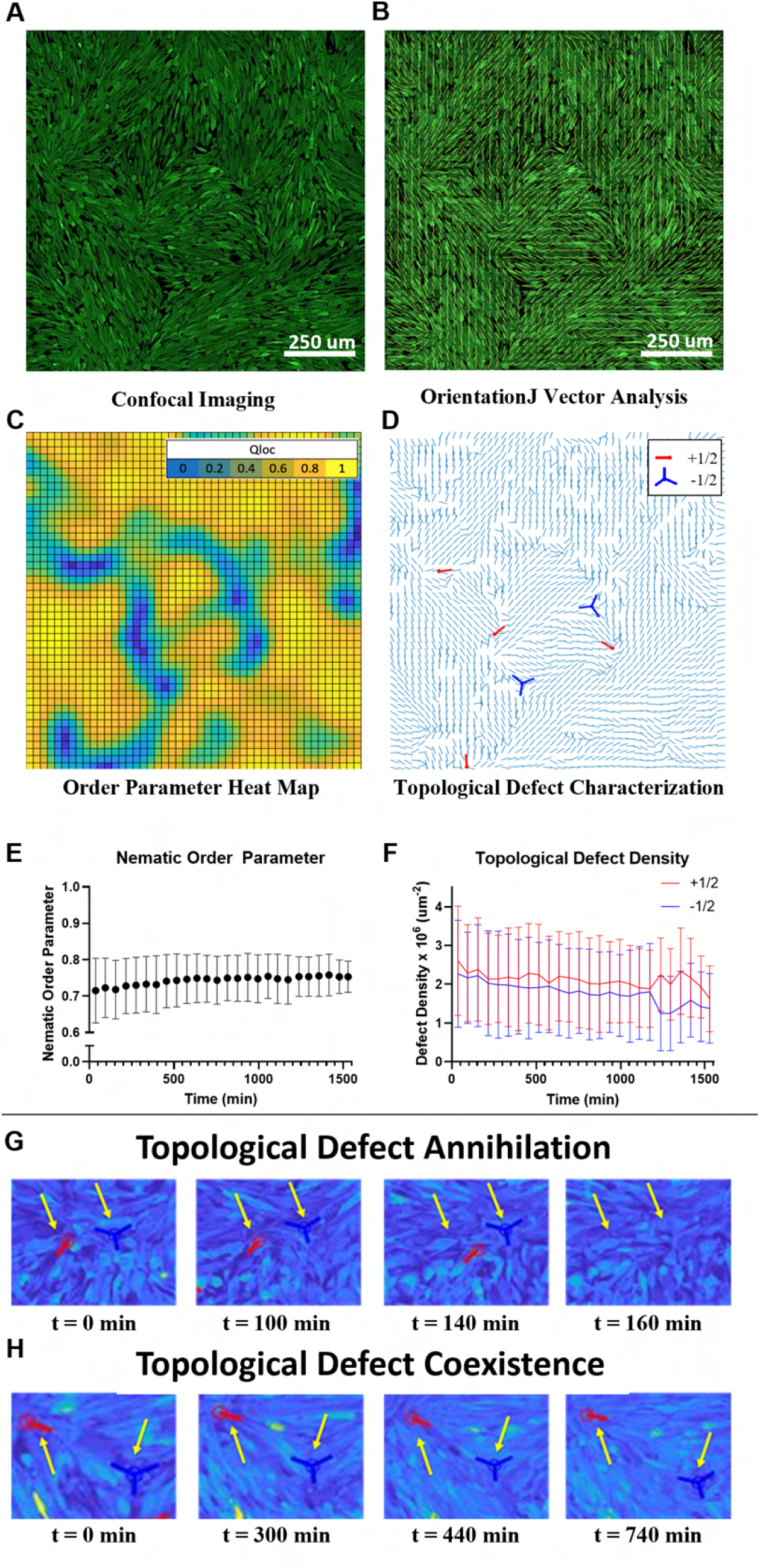
Dynamics of nematic order and topological defects in glioma cell cultures. **(A)** NPA glioma cells (cytosol tagged with green fluorescent protein (GFP)) imaged with confocal microscopy. **(B)** 2D director field (yellow dashes) for the image in panel A. **(C)** Heatmap of local order parameter, *Qloc* with coarse-graining length equal to 25 *μ*m for the same image in panel A. *Qloc* = 0 (dark blue) represents random alignment and *Qloc* = 1 (bright yellow) represents perfect nematic alignment. **(D)** Identification of topological defects with −1/2 defects (trefoils) in dark blue and +1/2 (comets) in red for the same image in panel A. Light blue dashes show the 2D director field. **(E)** Time evolution of the averaged *Qloc*. **(F)** Time evolution of the averaged topological defect density for −1/2(dark blue) and +1/2 (red) defects. **(G)** An example of the annihilation of a pair of topological defects with opposite charge. **(H)** An example of the extended coexistence of a pair of topological defects with opposite charge. For A,B: scale bars = 250 *μ*m. For E,F: *n* = 6 movies, *N* = 40 imaged positions; error bars correspond to SD; and time *t* = 0 min corresponds to one day post cell seeding. For G, H: scale bars = 10 *μ*m and *t* = 0 min was set arbitrarily.

**Figure S17:**
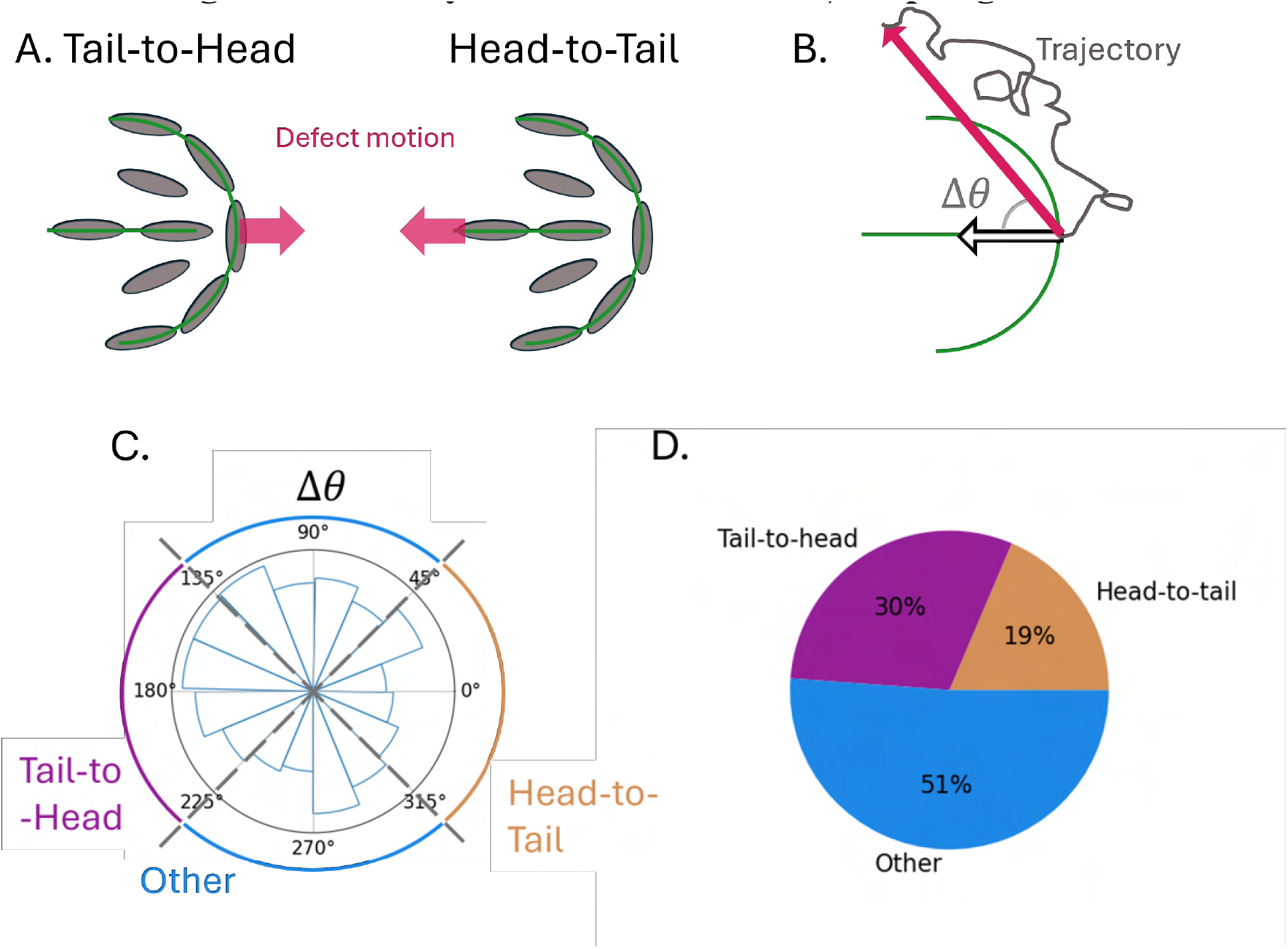
Analysis of the motion of +1/2 topological defects. **(A)** Two schematics of +1/2 topological defects (director field outline in green curves). Cells are colored in gray. The magenta arrows indicate the direction of defect motion for a tail-to-head moving defects (left panel) or a head-to-tail moving defect (right panel). **(B)** Schematic of a +1/2 topological defect with director field outlined in green. The gray curve represents the defect trajectory and the magenta arrow the displacement vector from the initial time point to the last time point. The empty arrow is the polarization **p** of the topological defect. In practice, the polarization vector was computed as the mean polarization vector over the trajectory. The angular difference between the mean polarization and the displacement vector is Δ*θ*. **(C)** Polar histogram of Δ*θ* for *N* = 172 independent +1/2 topological defects. The trajectories are classified as ‘head-to-tail’, if −45° < Δ*θ* < 45° (orange range), ‘tail-to-head’, if 225° < Δ*θ* < 135° (purple range), and ‘other’ in any other case (blue range). **(D)** Fraction of +1/2 topological defect with a motion head-to-tail, tail-to-head, or other.

## Notes

https://doi.org/10.5281/zenodo.15199294

### Competing Interest Statement

The authors have declared no competing interest.

### Summary of Updates

title and abstract slightly changed, and Figure 2 revised.

